# Dense, high-resolution mapping of cells and tissues from pathology images for the interpretable prediction of molecular phenotypes in cancer

**DOI:** 10.1101/2020.08.02.233197

**Authors:** James A. Diao, Wan Fung Chui, Jason K. Wang, Richard N. Mitchell, Sudha K. Rao, Murray B. Resnick, Abhik Lahiri, Chirag Maheshwari, Benjamin Glass, Victoria Mountain, Jennifer K. Kerner, Michael C. Montalto, Aditya Khosla, Ilan N. Wapinski, Andrew H. Beck, Amaro Taylor-Weiner, Hunter L. Elliott

**Author notes:** These authors contributed equally (James A. Diao, Wan Fung Chui, Jason K. Wang, Andrew H. Beck, Amaro Taylor-Weiner, Hunter L. Elliott).

## Abstract

While computational methods have made substantial progress in improving the accuracy and throughput of pathology workflows for diagnostic, prognostic, and genomic prediction, lack of interpretability remains a significant barrier to clinical integration. In this study, we present a novel approach for predicting clinically-relevant molecular phenotypes from histopathology whole-slide images (WSIs) using human-interpretable image features (HIFs). Our method leverages >1.6 million annotations from board-certified pathologists across >5,700 WSIs to train deep learning models for high-resolution tissue classification and cell detection across entire WSIs in five cancer types. Combining cell- and tissue-type models enables computation of 607 HIFs that comprehensively capture specific and biologically-relevant characteristics of multiple tumors. We demonstrate that these HIFs correlate with well-known markers of the tumor microenvironment (TME) and can predict diverse molecular signatures, including immune checkpoint protein expression and homologous recombination deficiency (HRD). Our HIF-based approach provides a novel, quantitative, and interpretable window into the composition and spatial architecture of the TME.

## Introduction

While manual microscopic inspection of histopathology slides remains the gold standard for evaluating the malignancy, subtype, and treatment options for cancer^1^, pathologists and oncologists increasingly rely on molecular assays to guide personalization of cancer therapy^2^. These assays can be expensive and time-consuming^3^, and unlike histopathology images, have not been historically and routinely collected, limiting their use in retrospective and exploratory research. Manual histological evaluation, on the other hand, presents several clinical challenges. Careful inspection requires significant time investment by board-certified anatomic pathologists and is often insufficient for prognostic prediction. Several evaluative tasks, including diagnostic classification, have also reported low inter-rater agreement across experts and low intra-rater agreement across multiple reads by the same expert^4,5^.

Modern computer vision methods present the potential for rapid, reproducible, and cost-effective clinical and molecular predictions. Over the past decade, the quantity and resolution of digitized histology slides has dramatically improved^6^. At the same time, the field of computer vision has made significant strides in pathology image analysis, including automated prediction of tumor grade^7^, mutational subtypes^8^, and gene expression signatures across cancer types^9,10^. In addition to achieving diagnostic sensitivity and specificity metrics that match or exceed those of human pathologists^11,12^, automated computational pathology can also scale to service resource-constrained settings where few pathologists are available. As a result, there may be opportunities to integrate these technologies into the clinical workflows of developing countries^13^.

However, end-to-end deep learning models that infer outputs directly from raw images present significant risks for clinical settings, including fragility of machine learning models to dataset shift, adversarial attack, and systematic biases present in training data^14–16^. Many of these risks stem from the well-known problem of model interpretability^17,18^. “Black-box” model predictions are difficult for users to interrogate and understand, leading to user distrust. Without reliable means for understanding when and how vulnerabilities may become failures, computational methods may face difficulty achieving widespread adoption in clinical settings^19,20^.

One emerging solution has been the automated computation of human-interpretable image features (HIFs) to predict clinical outcomes. HIF-based prediction models often mirror the pathology workflow of searching for distinctive, stage-defining features under a microscope and offer opportunities for pathologists to validate intermediate steps and identify failure points. In addition, HIF-based solutions enable incorporation of histological knowledge and expert pixel-level annotations which increases predictive power. Studied HIFs span a wide range of visual features, including stromal morphological structures^21^, cell and nucleus morphologies^22^, shapes and sizes of tumor regions^23^, tissue texture^24^, and the spatial distributions of tumor-infiltrating lymphocytes (TILs)^25,26^.

In recent years, the relationship between the TME and patient response to targeted therapies has been made increasingly clear^27,28^. For instance, immuno-supportive phenotypes, which exhibit greater baseline antitumor immunity and improved immunotherapy response, have been linked to the presence of TILs and elevated expression of programmed death-ligand 1 (PD-L1) on tumor-associated immune cells. In contrast, immuno-suppressive phenotypes have been linked to the presence of tumor-associated macrophages and fibroblasts, as well as reduced PD-L1 expression^28–30^. HIF-based approaches have the potential to provide an interpretable window into the composition and spatial architecture of the TME in a manner complementary to conventional genomic approaches. While prior HIF-based studies have identified many useful feature classes, most have been limited in scope. Studies to date often involve a single cell or tissue type; none have explored features that combine both cell and tissue properties. In addition, the majority of reported HIFs have only been vetted on a single cancer type, often non-small-cell lung cancer (NSCLC).

In this research study, we present a computational pathology pipeline that can integrate high-resolution cell- and tissue-level information from WSIs to predict treatment-relevant, molecularly-derived phenotypes across five different cancer types. In doing so, we introduce a diverse collection of 607 HIFs ranging from simple cell (e.g. density of lymphocytes in cancer tissue) and tissue quantities (e.g. area of necrotic tissue) to complex spatial features capturing tissue architecture, tissue morphology, and cell-cell proximity. Notably, we demonstrate that such features can generalize across cancer types and provide a quantitative and interpretable link to specific and biologically-relevant characteristics of each TME.

## Results

### Dataset characteristics and fully-automated pipeline design

In order to test our approach on a diverse array of histopathology images, we obtained 2,917 hematoxylin and eosin (H&E) stained, formalin-fixed and paraffin-embedded (FFPE) WSIs from the The Cancer Genome Atlas (TCGA), corresponding to 2,634 distinct patients. These images, each scanned at either 20x or 40x magnification, represented patients with skin cutaneous melanoma (SKCM), stomach adenocarcinoma (STAD), breast cancer (BRCA), lung adenocarcinoma (LUAD), and lung squamous cell carcinoma (LUSC) from 95 distinct clinical sites. We summarize the characteristics of TCGA patients in Supplemental Table 1. To supplement the TCGA analysis cohort, we obtained 4,158 additional WSIs for the five cancer types to improve model robustness.

To maximize capture of this information, we excluded images (n =91/2,917, 3.1%) if they failed basic quality control checks as determined by expert pathologists. Criteria for quality control were limited to mislabeling of tissue, excessive blur, or insufficient staining. For both TCGA and additional WSIs, we collected cell- and tissue-level annotations from a network of pathologists, amounting to >1.4 million cell-type point annotations and >200 thousand tissue-type region annotations (Supplemental Table 2).

We used the resulting slides and annotations to design a fully automated pipeline to extract HIFs from these slides (summarized in Figure 1a). First, we trained deep learning models for cell detection (“cell-type models”) and tissue region segmentation (“tissue-type models”). Training and validation of models was conducted on a development set of 1,561 TCGA WSIs, supplemented by the 4,158 additional WSIs (n =5719) (Figure 1b). Next, we exhaustively generated cell- and tissue-type model predictions for 2,826 TCGA WSIs, which were then used to compute a diverse array of HIFs for each WSI. Finally, we trained classical linear machine learning models to predict treatment-relevant molecular expression phenotypes using these HIFs.

**Fig. 1.**
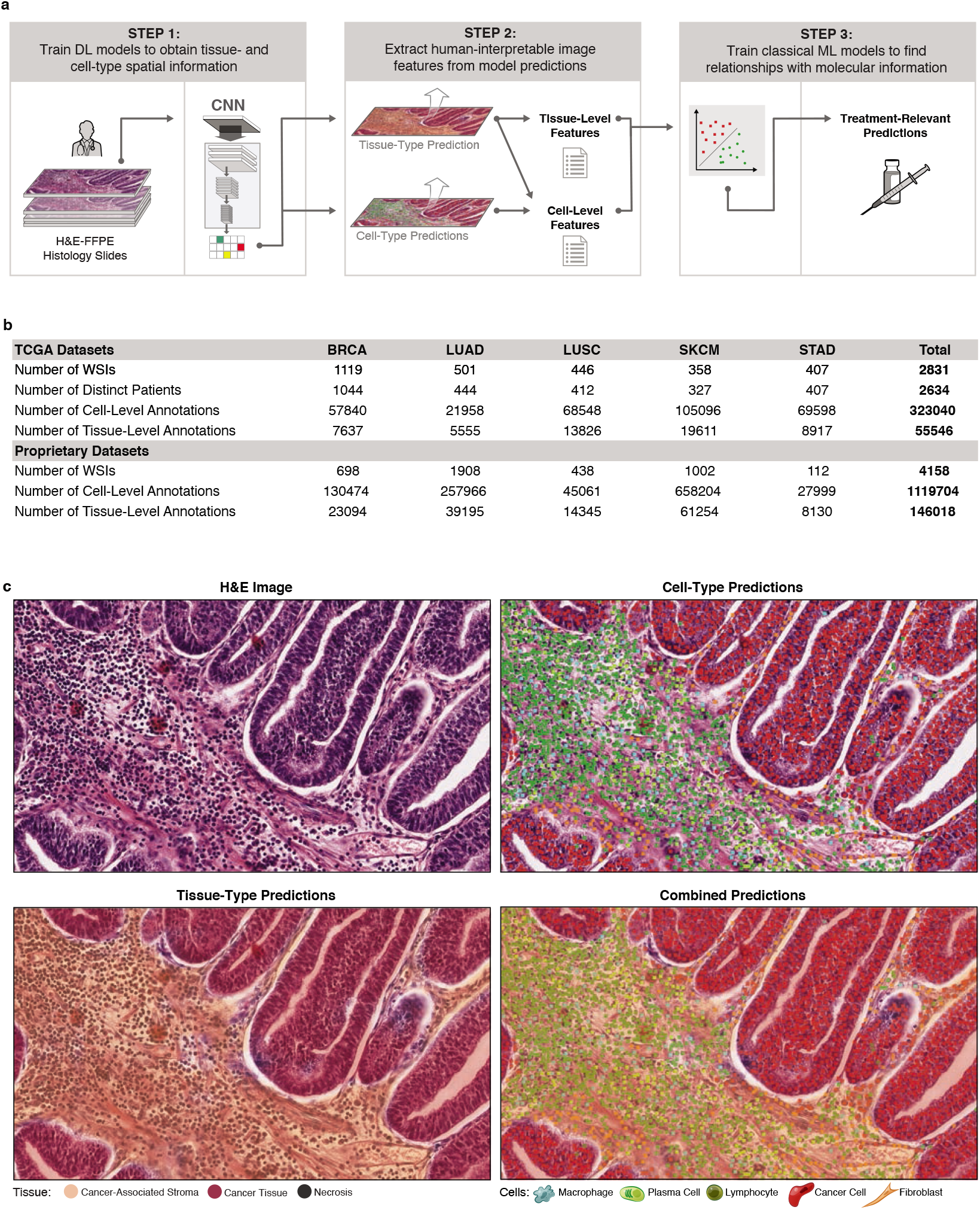
Dataset and pipeline overview. a) Methodology for extracting HIFs from high-resolution, digitized H&E images. b) Summary statistics on the number of WSIs, distinct patients, and annotations curated from TCGA and additional datasets. c) Unprocessed portions of STAD H&E-stained slides alongside corresponding heatmap visualizations of cell- and tissue-type predictions. Slide regions are classified into tissue types: cancer tissue (red), cancer-associated stroma (orange), necrosis (black), or normal (transparent). Pixels in cancer tissue or cancer-associated stroma areas are classified into cell types: lymphocyte (green), plasma cell (lime), fibroblast (orange), macrophage (aqua), cancer cell (red), or background (transparent).

### Cell- and tissue-type predictions yield a wide spectrum of HIFs

In the first step of our pipeline, we trained two convolutional neural networks (CNNs) per cancer type: (1) tissue-type models trained to segment cancer tissue, cancer-associated stroma, and necrotic tissue regions, and (2) celltype models trained to detect lymphocytes, plasma cells, fibroblasts, macrophages, and cancer cells. These models were improved iteratively through a series of quality control steps, including significant input from board-certified pathologists (Methods). These CNNs were then used to exhaustively generate cell-type labels and tissue-type segmentations for each WSI. We visualized these predictions as colored heatmaps projected onto the original WSIs (Figure 1c; Supplemental Figure 1). When quantified, these predictions capture broad multivariate information about the spatial distribution of cells and tissues in each slide.

Specifically, we used model predictions to extract 607 HIFs (Figure 2), which can be understood in terms of six categories (Figure 3). The first category includes cell type counts and densities across different tissue regions (e.g. density of plasma cells in cancer tissue) (Figure 3i-ii). The next category includes cell-level cluster features that capture intercellular spatial relationships, such as cluster dispersion, size, and extent (e.g. mean cluster size of fibroblasts in cancer-associated stroma) (Figure 3iii-iv). The third category captures cell-level proportion and proximity features, such as the proportional count of lymphocytes versus fibroblasts within 80 microns of the cancer-stroma interface (CSI) (Figure 3v-vi). The fourth category includes tissue area (e.g. mm^2^ of necrotic tissue) and multiplicity counts (e.g. number of significant regions of cancer tissue) (Figures Figure 3vii-viii). The fifth category includes tissue architecture features, such as the average solidity (“solidness”) of cancer tissue regions or the fractal dimension (geometrical complexity) of cancer-associated stroma (Figures Figure 3ix-x). The final category captures tissue-level morphology using metrics such as perimeter^2^ over area (shape roughness), lacunarity (“gappiness”), and eccentricity (Figure 3xi-xii). This broad enumeration of biologically-relevant HIFs explores a wide range of mechanisms underlying histopathology across diverse cancer types.

**Fig. 2.**
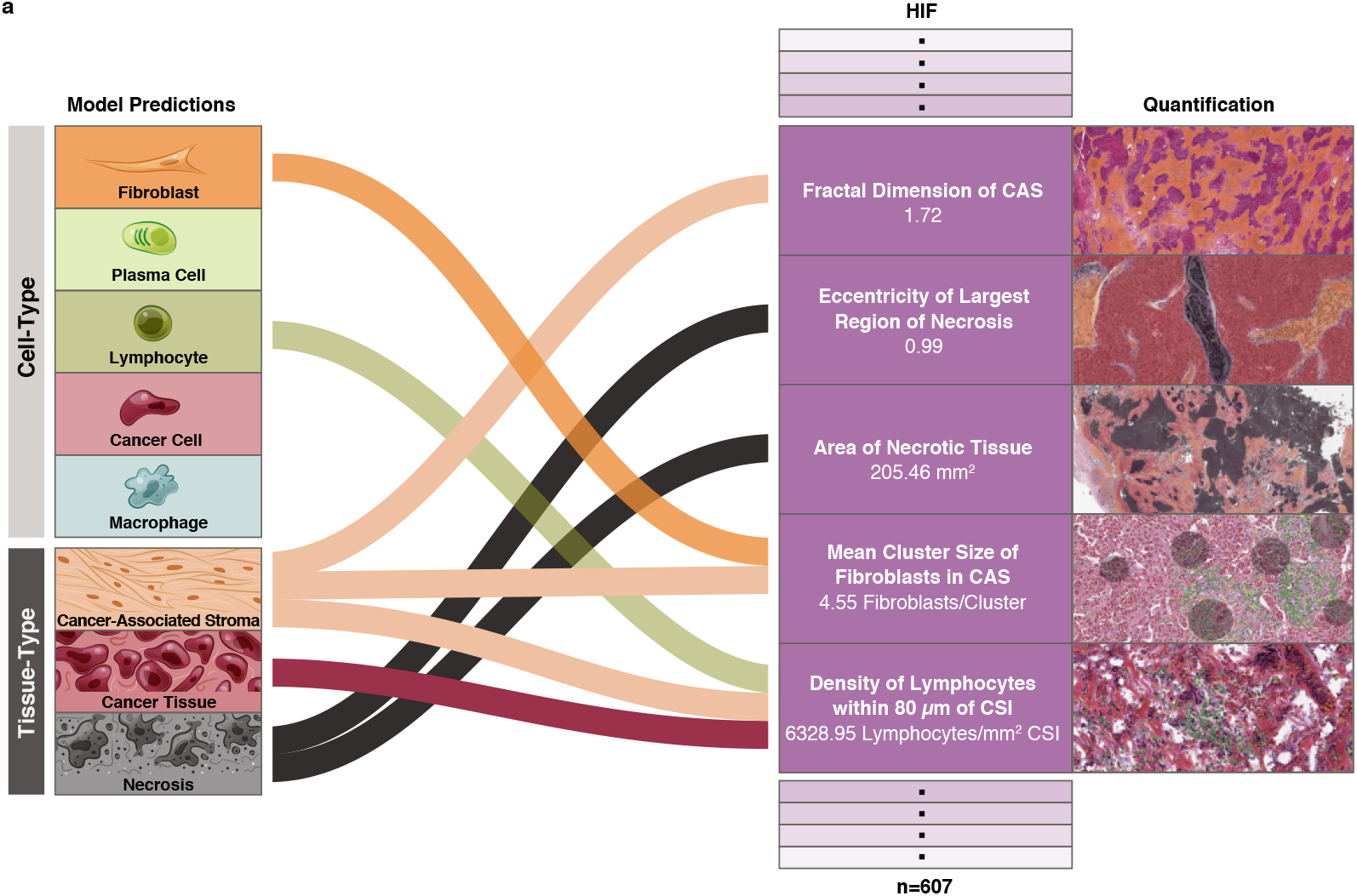
Human-interpretable image feature extraction workflow. Flow diagram of HIF extraction from model predictions for five example HIFs. For each HIF, an H&E snapshot with the corresponding cell- or tissue-type heatmap overlaid and the associated quantity are shown.

**Fig. 3.**
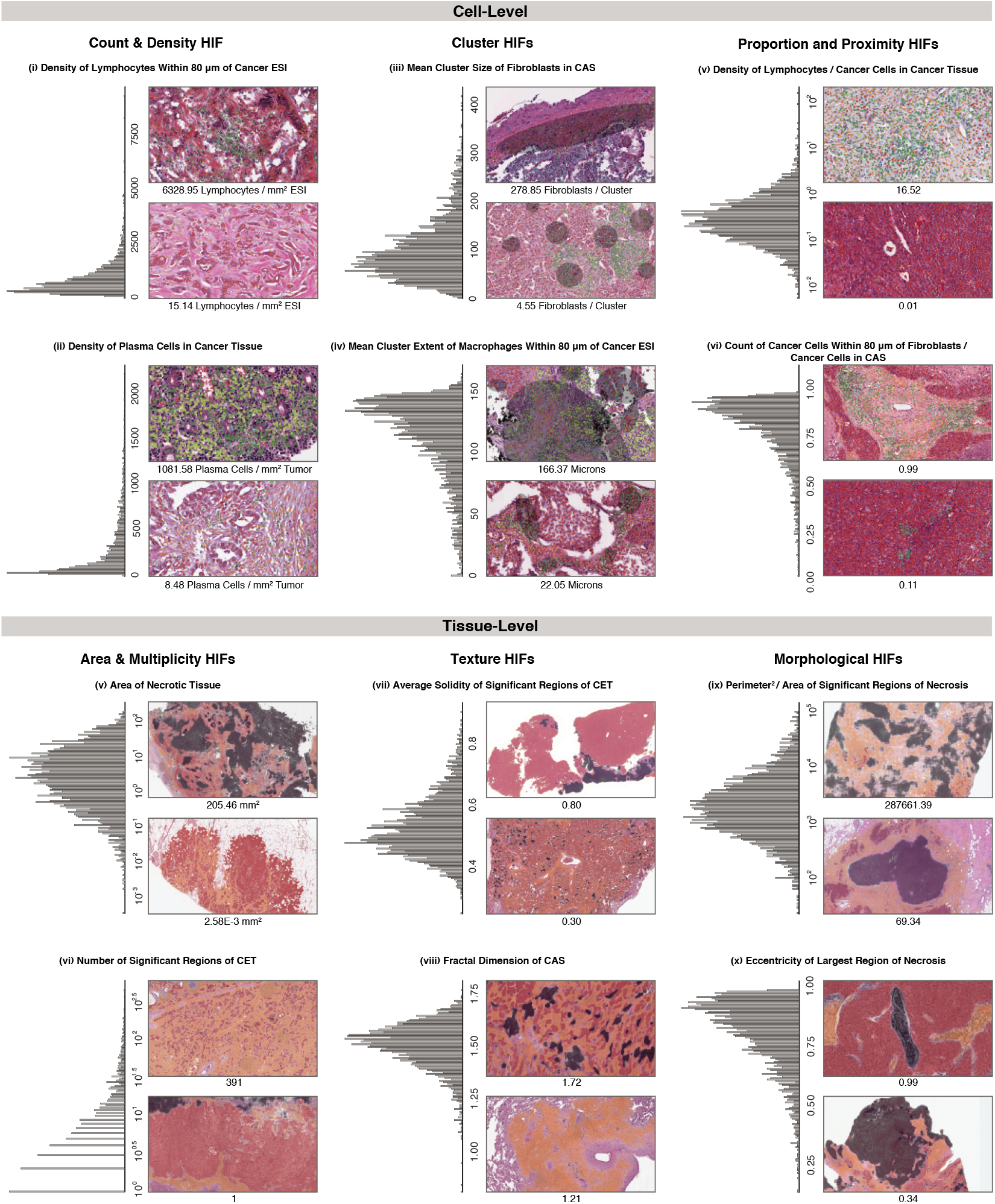
Overview of HIFs. Graphical overview of the 607 HIFs grouped into six categories: cell-level count and density (n = 56 HIFs), cell-level cluster (n = 180), cell-level proportion and proximity (n = 208), tissue-level area and multiplicity (n = 13), tissue-level architecture (n = 25), and tissue-level morphology (n = 125). For each HIF, a histogram of the HIF quantified in all patient samples across the five cancer types and H&E snapshots corresponding to high and low values with the corresponding heatmap are shown. Both snapshots are taken from patient samples of the same cancer type. Cell- and tissue-type heatmaps adhere to the same color scheme described in Figure 1c. In (iii), fibroblast clusters are annotated, contrasting one large cluster against multiple smaller clusters. In (iv), macrophage clusters and extents are annotated. Cluster extent is defined as the maximum distance between a cluster exemplar (defined via Birch clustering) and a cell within that cluster. Significant regions (viii) are defined as connected components (identified at the pixel-level) of a given tissue type with at least 10% the size of the largest connected component in the slide. A solidity (ix) of one corresponds to a completely filled object, while values less than one correspond to objects containing holes or with irregular boundaries. Fractal dimension (x) can efficiently estimate the geometrical complexity and irregularity of shapes and patterns, thus capturing tissue architecture. A fractal dimension of one corresponds to a perfectly smooth tissue border, while higher fractal dimension corresponds to increasing roughness and irregularity, indicating more extensive physical contact between adjacent tissue types. The fractal dimension of the CSI has been previously associated with dysfunction in antigen presentation^26^. Perimeter^2^ / Area (xi) is a unitless measure of shape roughness (e.g. square = 16, circle = 4π). Across all HIFs, tumor regions include cancer tissue (CT), cancer-associated stroma (CAS), and a combined CT+CAS.

### HIFs capture sufficient information to stratify cancer types

To visualize the global structure of the HIF feature matrix, we used Uniform Manifold Approximation and Projection (UMAP) to reduce the 607-dimensional HIF space into two dimensions (Figure 4a). The 2-D manifold projection of HIFs was able to separate BRCA, SKCM, and STAD into distinct clusters, while merging NSCLC subtypes LUAD and LUSC into one overlapping cluster (V-measure score = 0.47 using k-means with k = 4).

**Fig. 4.**
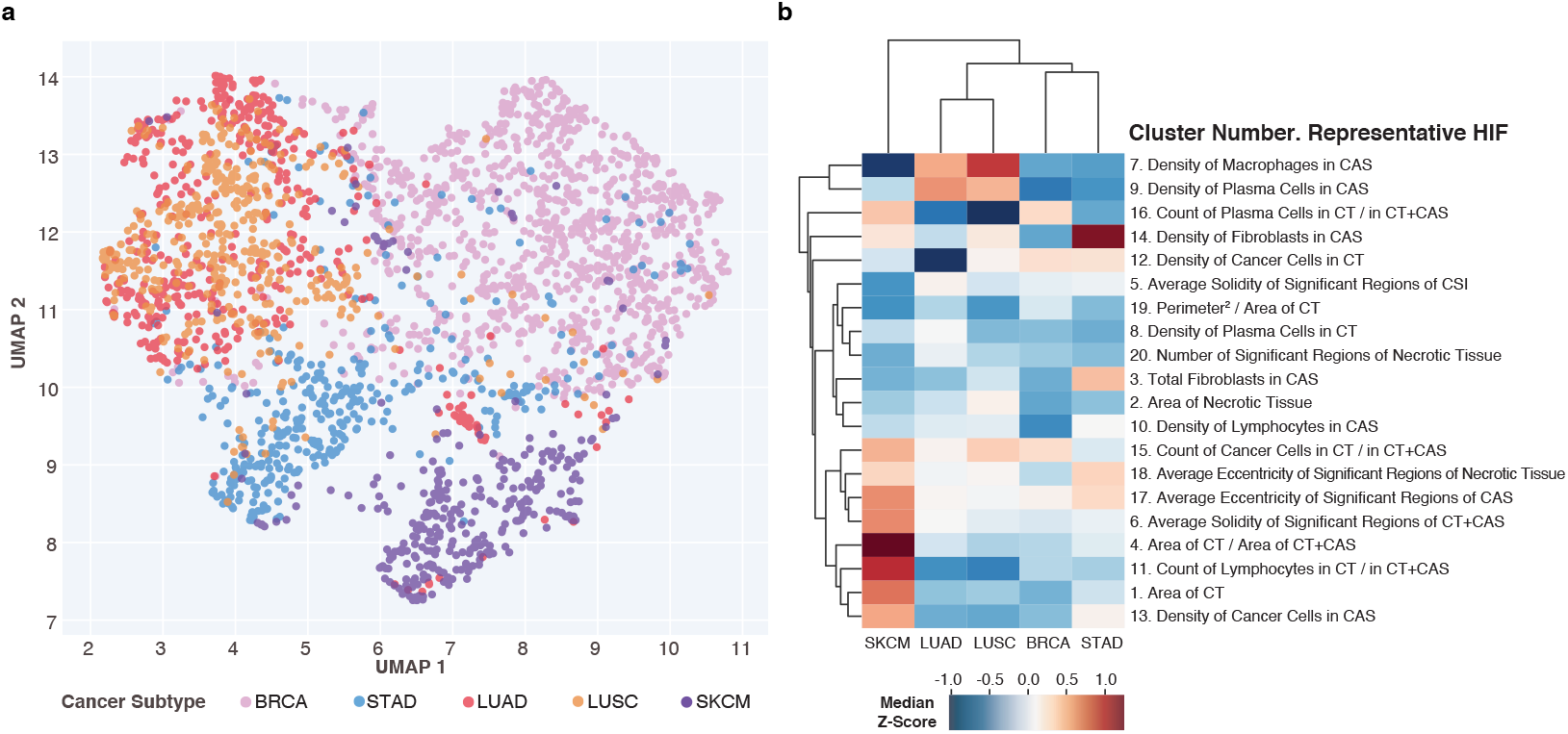
HIF differences across cancer types. a) UMAP projection and visualization of five cancer types reduced from the 607-dimension HIF space into two dimensions. Each point represents a patient sample colored by cancer type. b) Clustered heatmap of median Z-scores (computed pan-cancer) across cancer types for twenty HIFs, each representing one HIF cluster (defined pan-cancer). Hierarchical clustering was performed using average linkage and Euclidean distance. Clusters are annotated with a representative HIF chosen based on interpretability and high variance across cancer types.

Cancer type differences could be traced to specific and interpretable cell- and tissue-level features within the TME (Figure 4b). SKCM samples exhibited higher densities of cancer cells in cancer-associated stroma (pan-cancer median Z-score = 0.55, *P* < 10^−30^) and greater cancer tissue area per slide (Z-score = 0.72, *P* < 10^−30^) relative to other cancer types. These findings reflect biopsy protocols for SKCM, in which the excised region involves predominantly cancer tissue and minimal normal tissue. NSCLC subtypes LUAD and LUSC exhibited higher densities of macrophages in cancer-associated stroma (Z-score = 0.54 and 0.91, respectively; *P* < 10^−30^), reflecting the large population of macrophages infiltrating alveolar and interstitial compartments during lung inflammation^31^. NSCLC subtypes also exhibited higher densities of plasma cells (Z-score = 0.61 and 0.49; *P* < 10^−30^) in cancer-associated stroma, in agreement with prior findings in which proliferating B cells were observed in 35% of lung cancers^32,33^. STAD exhibited the highest density of lymphocytes in cancer-associated stroma (Z-score = 0.11, *P* = 2.16 × 10^−19^), corroborating prior work which identified STAD as having the largest fraction of TIL-positive patches per WSI among thirteen TCGA cancer types, including the five examined here^25^. Notably, HIFs are able to stratify cancer types by known histological differences without explicit tuning for cancer type detection.

### HIFs are concordant with sequencing-based cell and immune marker quantifications

To further validate our deep learning-based cell quantifications, we compared the abundance of the same cell type predicted by our cell-type models with those based on RNA sequencing^34^. Imagebased cell quantifications were correlated with sequencing-based quantifications across all patient samples and cancer types (pan-cancer) in three cell types (Supplemental Figure 2): leukocyte fraction (Spearman correlation coefficient (*ρ*) = 0.55, *P* < 2.2 × 10^−16^), lymphocyte fraction (*ρ* = 0.42, *P* < 2.2 × 10^−16^), and plasma cell fraction (*ρ* = 0.40, *P* < 2.2 × 10^−16^). Notably, imperfect correlation is expected as tissue samples used for RNA sequencing and histology imaging are extracted from different portions of the patient’s tumor, and thus vary in TME due to spatial heterogeneity.

There is significant correlation structure among individual HIFs due to the modular process by which feature sets are generated, as well as inherent correlations in underlying biological phenomena. For example, proportion, density, and spatial features of a given cell or tissue type all rely on the same underlying model predictions. In order to identify mechanistically-relevant and inter-correlated groups of HIFs, hierarchical agglomerative clustering was conducted (Methods; Supplemental Data 1). This clustering also increases the power of multiple-hypothesis-testing corrections by accounting for feature correlation^35^. Pan-cancer HIF clusters strongly correlated with immune markers of leukocyte infiltration, IgG expression, TGF-*β* expression, and wound healing (Figure 5a), each quantified by scoring bulk RNA sequencing reads for known immune expression signatures. We conducted the same correlational analysis for each cancer type individually, and observed high concordance among the top-correlated HIF clusters per immune marker (Supplemental Table 3).

**Fig. 5.**
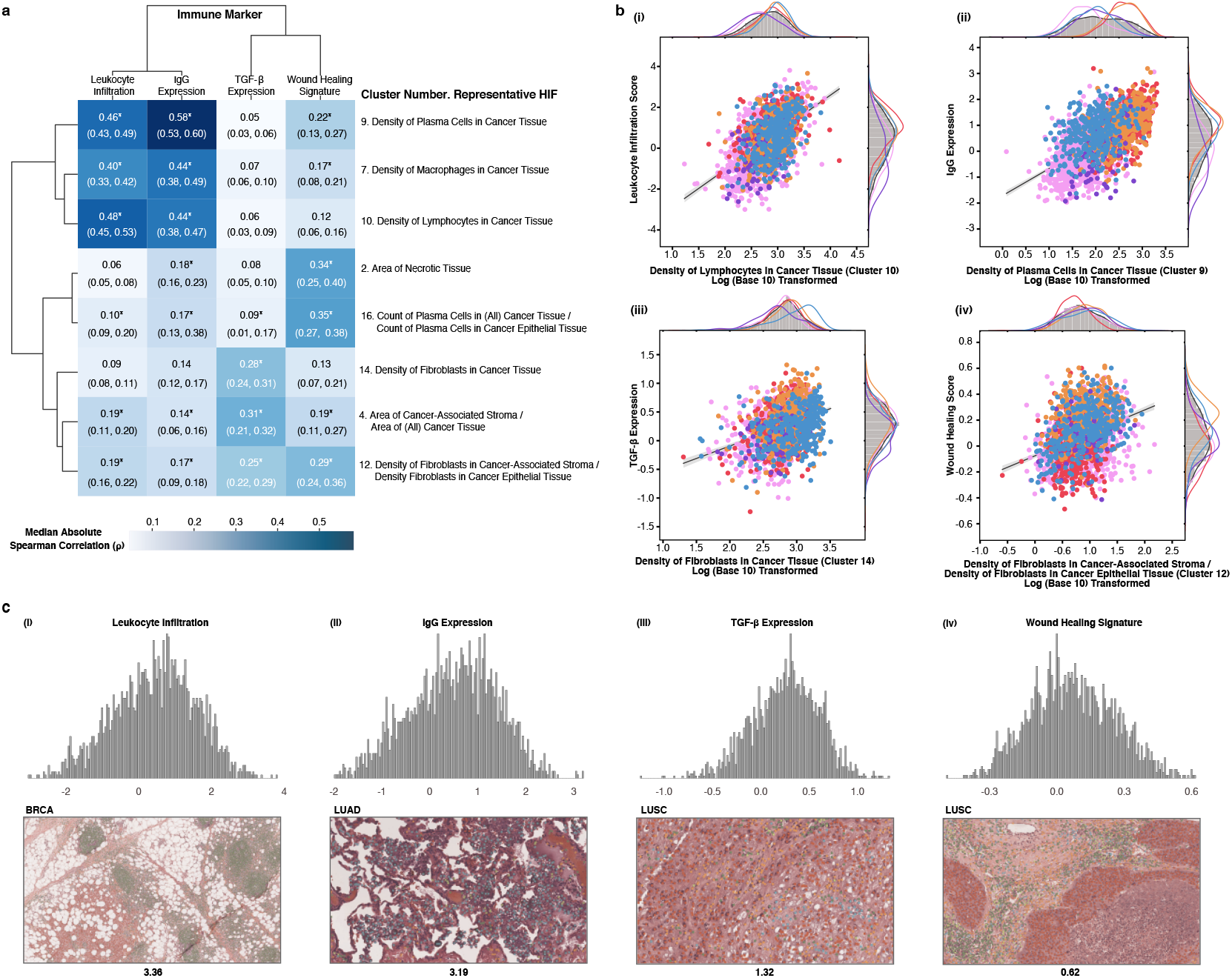
Validation of HIFs against immune markers. a) Clustered heatmap of median absolute Spearman correlation coefficients (*ρ*) computed across all patient samples between eight HIF clusters (defined pan-cancer) and four canonical immune markers. Hierarchical clustering was done using average linkage and euclidean distance. Median absolute Spearman correlation coefficients with a combined (via the Empirical Brown’s method) and corrected (via the Benjamini-Hochberg procedure) P value lower than the machine precision level (10^−30^) are annotated with an asterisk. Negative control analyses are included in Supplemental Table 3. Tumor regions include cancer tissue (CT), cancer-associated stroma (CAS), and a combined CT+CAS. b) Correlation and kernel density estimation plots between representative HIFs and immune markers. Points are colored by cancer type. X-axes are log-transformed (base ten). Trendlines are plotted on the log-transformed data. Cell densities are reported in count/mm^2^ and tissue areas are reported in mm^2^. c) Histogram of immune marker expression (Z-score) across all patients, alongside an H&E snapshot with its cell-type heatmap overlaid corresponding to high expression of the given immune marker. Cell-type heatmaps adhere to the same color scheme described in Figure 1c.

Molecular quantification of leukocyte infiltration was concordant with the density of leukocyte-lineage cells in cancer tissue plus cancer-associated stroma (CT+CAS) quantified by our deep learning pipeline, including lymphocytes (median absolute Spearman correlation *p* for associated HIF cluster= 0.48, *P* < 10^−30^; Figure 5bi), plasma cells (cluster *p* = 0.46, *P* < 10^−30^), and macrophages (cluster *p* = 0.40, *P* < 10^−30^). Similarly, we observed associations between IgG expression and the density of leukocyte-lineage cells in CT+CAS, with plasma cells being the most strongly correlated (cluster *p* = 0.58, *P* < 10^−30^), as expected given their role in producing immunoglobulins (Figure 5bii). TGF-*β* expression was associated with the density of fibroblasts in CT+CAS (cluster *ρ* = 0.28, *P* < 10^−30^; Figure 5biii), building upon prior studies which found that TGF-*β*1 can promote fibroblast proliferation^36–38^. TGF-*β* expression was also correlated with the area of cancer-associated stroma relative to CT+CAS (cluster *ρ* = 0.31, *P* < 10^−30^), shedding further light on the role of stromal proteins in modulating TGF-*β* levels^39^. Wound healing signature was positively associated with the density of fibroblasts in cancer-associated stroma versus in cancer tissue (cluster *ρ* = 0.29, *P* < 10^−30^; Figure 5biv), which corroborates findings that both tumors and healing wounds alike modulate fibroblast recruitment and proliferation to facilitate extracellular matrix deposition^40^. H&E snapshots corresponding to high expression of each of the four immune markers are shown in Figure 5c with corresponding cell-type heatmaps overlaid.

### HIFs are predictive of clinically-relevant phenotypes

To evaluate the capability of HIFs to predict expression of clinically-relevant, immuno-modulatory genes, we conducted supervised prediction of binarized classes for five clinically-relevant phenotypes: (1) programmed cell death protein 1 (PD-1) expression, (2) PD-L1 expression, (3) cytotoxic T-lymphocyte-associated protein 4 (CTLA-4) expression, (4) HRD score, and (5) T cell immunoreceptor with Ig and ITIM domains (TIGIT) expression (Figure 6; Supplemental Figure 3). Using the 607 HIFs computed per WSI, predictions were conducted for cancer types individually as well as pan-cancer. SKCM predictions were conducted only for TIGIT expression due to insufficient sample sizes for the remainder of outcomes (Methods). To demonstrate model generalizability across varying patient demographics and sample collection processes, area under the receiver operating characteristic (AUROC) and area under the precision-recall curve (AUPRC) performance metrics were computed on hold-out sets composed exclusively of patient samples derived from tissue source sites not seen in the training sets (Supplemental Table 4).

**Fig. 6.**
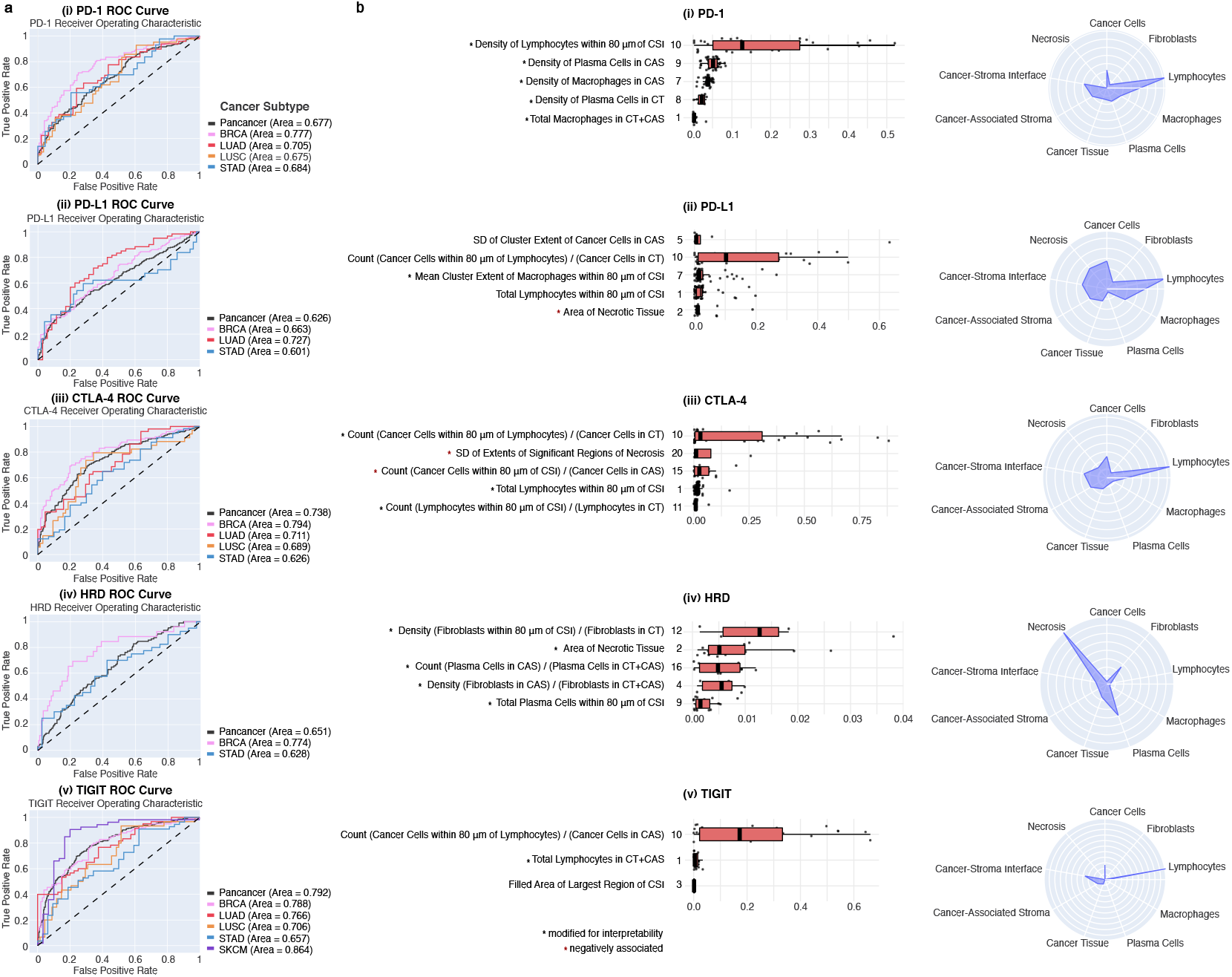
HIF-based prediction of molecular phenotypes. a) ROC curves for (i) PD-1, (ii) PD-L1, (iii) CTLA-4, (iv) HRD, and (v) TIGIT hold-out predictions across cancer types and pan-cancer. SKCM predictions were conducted only for TIGIT due to low sample sizes. Pan-cancer predictions use binary labels thresholded independently by cancer type. For TIGIT predictions, pan-cancer includes all five cancer types. For the remainder of predictions, pan-cancer includes all cancer types excluding SKCM. Random classifiers correspond to AUROC = 0.50. b) Visualization of predictive HIFs for each molecular phenotype. Boxplots show the top five most predictive HIF clusters for each phenotype in pan-cancer models. For TIGIT predictions, pan-cancer models only included three non-zero HIF clusters. Clusters are ranked by the maximum absolute ensemble beta across HIFs in a given cluster. Ensemble betas are computed per HIF as the average across the three models incorporated into the final ensemble evaluated on the hold-out set. Each boxplot highlights the median and interquartile range for ensemble betas in each cluster. Each cluster is labeled with a representative HIF corresponding to the maximum absolute ensemble beta value. In cases where that HIF is difficult to interpret, a more interpretable HIF within a five-fold difference of the maximum ensemble beta is presented (indicated by a black asterisk). As absolute values were used for ranking, HIFs with negative ensemble betas are denoted by a red asterisk. Boxplots of predictive HIF clusters forcancertype-specific models are included in Supplemental Figure 5. Radar charts show the normalized magnitude of ensemble betas in pan-cancer models stratified across nine HIF axes, corresponding to the five cell types, three tissue types, and CSI. Normalized magnitudes were computed as the sum of absolute ensemble betas for HIFs associated with each axis divided by the total number of HIFs associated with said axis (e.g. all HIFs involving fibroblasts). Multiple predictive HIFs are visualized with overlaid cell- or tissue-type heatmaps in Figure 3. Tumor regions include cancer tissue (CT), cancer-associated stroma (CAS), and a combined CT+CAS.

HIF-based models were not predictive for every phenotype in each cancer type (hold-out AUROC < 0.6; see Supplemental Table 5 for all results including negatives). In the successful prediction models (hold-out AUROC range = 0.6010.864; Figure 6a), precision-recall curves revealed that models were robust to class-imbalance, achieving AUPRC performance surpassing positive class prevalence by 0.104-0.306 (Supplemental Figure 4). Notably, AUROC performance of our HIF-based linear model for PD-L1 expression in LUAD was comparable to that achieved by “black-box” deep learning models trained on hundreds of thousands of paired H&E and PD-L1 example patches in NSCLC^41^.

### Predictive HIFs provide interpretable link to clinically-relevant phenotypes

Interpretable features enable interrogation and further validation of model parameters as well as generation of new biological hypotheses. Towards this end, for each prediction task we identified the five most important HIF clusters as determined by magnitude of model coefficients (Figure 6b; Supplemental Figure 5) and computed cluster-level P-values to evaluate significance (Supplemental Table 6; Methods).

As expected, prediction of PD-1 and PD-L1 involved similar HIF clusters (Pearson correlation between PD-1 and PD-L1 expression = 0.53; Supplemental Figure 6). The count of cancer cells within 80 microns of lymphocytes, as well as the density of lymphocytes in CT+CAS, was significantly selected during model fitting for both of PD-1 and PD-L1 expression in pan-cancer and BRCA models (Figure 6bi-ii; Supplemental Figures 5i-ii). Furthermore, in both LUAD and LUSC, the count of lymphocytes in CT+CAS was similarly predictive of PD-1 and PD-L1 expression. The importance of these HIFs which capture lymphocyte infiltration between and surrounding cancer cells corroborates prior literature which demonstrated that TILs correlated strongly with higher expression levels of PD-1 and PD-L1 in early breast cancer42 and NSCLC^43,44^.

The area, morphology, or multiplicity of necrotic tissue proved predictive of PD-1 expression in LUAD, LUSC, and STAD models and of PD-L1 expression in pan-cancer, BRCA, and LUAD models, expanding upon prior findings that tumor necrosis correlated positively with PD-1 and PD-L1 expression in LUAD^45^. The density, proximity, or clustering properties of plasma cells was predictive of PD-1 expression in all models excluding LUAD, suggesting a role for plasma cells in modulating PD-1 expression. Recent studies in SKCM, renal cell carcinoma, and soft-tissue sarcoma have demonstrated that an enrichment of B-cells in tertiary lymphoid structures was positively predictive of response to immune checkpoint blockade therapy^46–48^. The density of fibroblasts in cancer-associated stroma or within 80 microns of the CSI was predictive of PD-L1 expression in LUAD and STAD, respectively, corroborating earlier discoveries that cancer-associated fibroblasts promote PD-L1 expression^49^.

Less is known about the relationship between the TME and CTLA-4 expression. By investigating predictive HIFs we can begin to enumerate features of the TME that correlate with CTLA-4 expression. The proximity of lymphocytes to cancer cells (pan-cancer and BRCA), morphology of necrotic regions (LUAD and LUSC), and density of cancer cells in CT+CAS versus exclusively in cancer-associated stroma (BRCA and STAD) were predictive of CTLA-4 expression across multiple models (Figure 6biii; Supplemental Figure 5iii).

Area of necrotic tissue (pan-cancer and BRCA) as well as various morphological properties of necrotic regions including perimeter and lacunarity (BRCA and STAD) were predictive of HRD (Figure 6biv; Supplemental 5iv). In HRD, ineffective DNA damage repair can result in the accumulation of severe DNA damage and subsequent cell death through apoptosis as well as necrosis^50,51^. The density and count of fibroblasts near or in cancer-associated stroma was also predictive of HRD in the pan-cancer and BRCA models, corroborating prior findings that persistent DNA damage and subsequent accumulation of unrepaired DNA strand breaks can induce reprogramming of normal fibroblasts into cancer-associated fibroblasts^52^.

Like the three other immune checkpoint proteins (PD-1, PD-L1, and CTLA-4), TIGIT expression was also associated with markers of tumor inflammation, including the count of cancer cells within 80 microns of lymphocytes (pan-cancer and BRCA), the total number of lymphocytes in CT+CAS (pan-cancer and BRCA), and the proportional count of lymphocytes to cancer cells within 80 microns of the CSI (LUAD) (Figure 6bv; Supplemental Figure 5v). These findings corroborate prior findings that TIGIT expression, alongside PD-1 and PD-L1 expression (Pearson correlation between TIGIT and PD-1 = 0.84; TIGIT and PD-L1 = 0.56; Supplemental Figure 6), is correlated with TILs^53^. HIF clusters capturing morphology and architecture of necrotic tissue (e.g. fractal dimension, lacunarity, extent, perimeter^2^ / area) were associated with TIGIT expression in LUAD, LUSC, SKCM, and STAD models, although these relationships have yet to be investigated.

## Discussion

Our study is the first to demonstrate the value of combining deep learning-based cell- and tissue-type classifications to compute image features that are both biologically-relevant and human-interpretable. We demonstrate that computed HIFs can recapitulate sequencing-based cell quantifications, capture canonical immune markers such as leukocyte infiltration and TGF-*β* expression, and robustly predict five molecular phenotypes relevant to oncology treatment efficacy and response. We also demonstrate the generalizability of our associations, as evidenced by similarly predictive HIF clusters across biopsy images derived from five different cancer types. While prior studies have applied deep learning methodologies to capture cell-level information, such as the spatial configuration of immune and stromal cells^26,54^, or tissue-level information55 alone, our combined cell plus tissue approach enables quantification of increasingly complex and expressive features of the TME, ranging from the mean cluster size of fibroblasts in cancer-associated stroma to the proximity of TILs or cancer-associated fibroblasts to the CSI. By training models to make six-class cell-type and four-class tissue type classifications, our approach is also able to aggregate more layers of information than prior studies. Indeed, while TILs are emerging as a promising biomarker in solid tumors such as triple-negative and HER2-positive breast cancer^54^, TILs differ from stromal lymphocytes, and substantial signal can be obtained by considering multiple cell-tissue combinations^21^.

Our approach of exhaustively generating cell- and tissue-type predictions across entire WSIs at subcellular resolution (of two and four microns, respectively) is novel to the best of our knowledge, and improves upon previous tiling approaches that downsample the image. The tissue visible in a WSI is already only a fraction of the tumor itself. Using the entire slide (rather than selected tiles) reduces the probability of fixating on non-generalizable local effects and enables quantification of complex characteristics that span multiple tissue regions (e.g. multiplicity, solidity, and fractal dimension of significant necrotic regions).

In addition, our approach of capturing specific and interpretable features of the tumor and its surroundings can facilitate hypothesis generation and enable a deeper understanding of the TME’s influence on drug response. Indeed, recent studies provide evidence that tumor immune architecture can greatly dictate clinical efficacy of immune checkpoint inhibitor56 and poly (ADP-ribose) polymerase (PARP) inhibitor therapies^57^.

Lastly, during both model development and evaluation, we sought to emphasize robustness to real-world variability^58^. In particular, we supplemented TCGA WSIs with additional diverse datasets during CNN training, integrated pathologist feedback into model iterations, and evaluated HIF-based model performance on hold-out sets composed exclusively of samples from unseen tissue source sites, improving upon prior approaches to predicting molecular outcomes from TCGA H&E images^22,59^.

One major limitation of machine-learning approaches is the quality of training data. While our cell and tissue classification models were trained on a combination of TCGA and additional datasets, molecular associations and predictions were derived solely from TCGA. Biopsy images submitted to the TCGA dataset suffer from selection bias towards more definitive diagnoses and early-stage disease that may not generalize well to ordinary clinical settings. Moreover, the images only contain H&E staining, which limits the amount of information available to us. It is possible that integrating multimodal data containing stains against Ki-67 or immunohistological targets may increase confidence in cell classifications^60^. In addition to the quality of slide images, annotations are also variable in reliability. Macrophages are particularly difficult for pathologists to identify solely under H&E staining. While the accuracy of an individual pathologist identifying macrophages may be poor, our models represent a consensus across hundreds of pathologist annotators which may carry a more reliable signal^61,62^.

Furthermore, morphologically similar cells (e.g. macrophages, dendritic cells, endothelial cells, pericytes, myeloid derived suppressor cells, and atypical lymphocytes) may all be captured under a single cell-type prediction. Thus, HIFs may, in reality, capture information about a mixture of cell types. For example, in diffuse forms of STAD in which cancer cells invade smooth muscle tissue, our models misclassified certain smooth muscle cells as fibroblasts. Therefore, fibroblast-label HIFs likely reflect a mixture of these two cell types in STAD. Further disambiguation of morphologically-similar cell types may decrease noise in HIF estimates and improve performance.

These interpretable sets of HIFs, computed from tens-of-thousands of deep learning-based cell- and tissue-type predictions per patient, improve upon conventional “black-box” approaches which apply deep learning directly to WSIs, yielding models with millions of parameters and limited interpretability. Recent work has revealed the weaknesses of low-interpretability models, including brittleness to dataset shift, vulnerabilities to adversarial attack, and susceptibility to the biases of the data-generative process. Unlike class activation maps utilized in prior studies as a heuristic to identify predictive image regions^9,10^, HIFs can be interpreted in aggregate across thousands of images and mapped directly onto biological concepts.

Beyond suggesting interpretable hypotheses for causal mechanisms (e.g. the anti-tumor effect of high lymphocyte density), our HIF-based approach can be continually validated at several points: pathologists can judge the quality of cell and tissue-type predictions, estimate the values of each relevant feature using traditional manual scoring, and observe whether there is a significant failure given real-world variability in sample preparation and quality. Unlike “black-box” models that may opaquely rely on features that are predictive but disconnected from the outcome of interest, such as tissue excision or preparation artifacts (e.g. surgical or pathologist markings)^16,19^, HIF-based predictions can be traced to observable features, allowing model failures to be explained and addressed. While performance is vitally important in clinical settings and additional studies comparing end-to-end and HIF-based approaches are needed, the improved trust and reliability against unexpected failures make HIF-based models a valuable alternative.

Finally, the ability to predict molecular phenotypes directly from WSIs in an interpretable fashion has numerous potential benefits for clinical oncology. Hospitals, healthcare institutions, and pharmaceutical and biotechnology companies have decades of archival histopathology data captured from routine care and clinical trials^63^. HIF-based models capable of capturing molecular information could supplement molecular assays that are often expensive and time-consuming^3^, enable the discovery of novel patient subpopulations with specific disease processes and treatment susceptibilities, and generate hypotheses for subsequent research.

## ACKNOWLEDGEMENTS

We are grateful to the software engineering and machine learning teams at PathAI, Inc. for developing the systems and pipelines used for model development and feature extraction. We also than k the Harvard-MIT Program in Health Sciences and Technology for enabling J.A.D., W.F.C., and J.K.W. to conduct this work towards satisfaction of their research theses. This work was funded by PathAI, Inc.

## AUTHOR CONTRIBUTIONS

J.A.D., W.F.C., J.K.W., A.T.W, H.L.E., and A.H.B. conceived the project. J.A.D. and W.F.C. trained the models, generated heatmaps, and computed image features. A.L. and C.M. assisted with model troubleshooting and feature computation. S.K.R. and M.B.R. provided feedback on predicted heatmaps to enable iterative model improvements. B.G. and I.N.W. supervised collection of pathologist annotations for model training. J.A.D. and J.K.W. conducted statistical analysis of image feature associations. J.K.W. developed models for molecular phenotype prediction and image feature visualization. H.L.E. and A.T.W. supervised model training and statistical analyses. All authors contributed to preparation of the manuscript.

## Methods

### Dense, high-resolution prediction of cell and tissue types using convolutional neural networks

In order to compute histopathological image features for each slide, it was necessary to first generate cell and tissue predictions per WSI. To this end, we asked a network of board-certified pathologists to label WSIs with both polygonal region annotations based on tissue type (cancer tissue, cancer-associated stroma, necrotic tissue, and normal tissue or background) and point annotations based on cell type (cancer cells, lymphocytes, macrophages, plasma cells, fibroblasts, and other cells or background). This collection of expert annotations was then used to train six-class cell type and four-class tissue-type classifiers.

Several steps were taken to ensure the accuracy and generalizability of our models. First, it was important to recognize that common cell and tissue types, such as cancer-associated stroma or cancer cells, show morphological differences between BRCA, LUAD, LUSC, SKCM, and STAD. As a result, we trained separate cell- and tissue-type detection models for each of these five cancer types, for a total of ten models. Second, it was important to ensure that our models did not overfit to the histological patterns found in the training set. To avoid this, we followed the conventional protocol of splitting our data into training, validation, and test sets, and incorporated additional annotations of the same five cancer types from PathAI’s databases into the model development process. Together, these datasets represented a wide diversity of examples for each class in each cancer type, thus improving the generalizability of these models beyond the TCGA dataset.

Using the combined dataset of annotated TCGA and additional WSIs, we trained deep convolutional neural networks (CNN) to output dense pixelwise cell- and tissue-type predictions at a subcellular spatial resolution of two and four microns, respectively (spatial resolution dictated by stride). To ensure that our models achieved sufficient accuracy for feature extraction, models were trained in an iterative process, with each updated model’s predictions visualized as heatmaps to be reviewed by board-certified pathologists. In heatmap visualizations, tissue categories were segmented into colored regions, while cell types were identified as colored squares. This process continued until there were minimal systematic errors and the pathologists deemed the model sufficiently trustworthy for feature extraction.

### Pathologist validation of cell- and tissue-type predictions

During the CNN training process, we worked iteratively with three board-certified pathologists to conduct subjective evaluation of model predictions to inform multiple rounds of training. CNN models were initially trained on a set of primary annotations collected from the pathologist network. Following the conclusion of each training round (defined by model convergence), predicted cell and tissue heatmaps were reviewed for systematic errors (e.g. overprediction of fibroblasts, macrophages, and plasma cells, underprediction of necrotic tissue). New annotations would then be collected from the pathologist network focusing on areas of improvement (e.g. mislabeled macrophages) to initiate a subsequent training round.

### Tissue-based feature extraction

Using the tissue-type predictions, we extracted 163 different region-based features from each WSI in the TCGA dataset. Each of these features belonged to one of three general categories.

The first category consisted of areas (n = 13 HIFs). By simple pixel summation, we computed the total areas (in mm^2^) of cancer tissue, cancer-associated stroma, cancer tissue plus cancer-associated stroma, regions at the cancer-stroma interface, and necrosis in each slide. These features are interpretable and technically attainable by human pathologists, but would be prohibitively time-consuming and inconsistent across pathologists to calculate in practice.

The second category, which contributed the bulk of the features, made use of the publicly available scikit-image.measure.regionprops module to find the connected components of each of these tissue types at the pixel-level using eight-connectivity. Once these connected components were found, we used both library-provided and self-implemented methods to extract a series of morphological features (n = 125 HIFs), similar to the approach suggested by Wang et al. in 2018^23^. These HIFs measured a wide variety of tissue characteristics, ranging from quantitative, size-based measures like the number of connected components, major and minor axis lengths, convex areas, and filled areas, to more qualitative, shape-based measures like Euler numbers, lacunarity, and eccentricity. Recognizing the log-distribution of connected component size, we computed these features not just across all connected components, but also for both the largest connected component only and across the most “significant” connected components, defined as components larger than 10% the size of the largest connected component. In aggregating metrics across considered components, we incorporated both averages and standard deviations of HIFs (e.g. standard deviation of eccentricities of significant regions of necrosis), to capture both summary metrics and metrics of intratumor heterogeneity.

The third category of features captures tissue architecture (n = 25 HIFs). Inspired by Lennon et al.^24^, we calculated the fractal dimensions and solidity measures of different tissue types, capturing both the roundness and filled-ness of the tissue, under the hypothesis that the ability for these measures to separate different subtypes of lung cancer might translate to a similar ability to predict clinically-relevant phenotypes. These features allowed us to capture information about how tissue filled up space, rather than just the summative sizes and shapes captured by the first and second categories.

### Cell- and tissue-based feature extraction

After obtaining six-class cell-type predictions for each pixel of a WSI, we generated five binary masks corresponding to each of the five specified cell types. We then combined cell- and tissue-level masks to compute properties of each cell type in each tissue type (e.g. fibroblasts in cancer-associated stroma), extracting 444 HIFs.

An initial group of features that were readily calculable from our model predictions included simple counts and densities of cell types in different tissue types. For example, an overlay of a particular slide’s lymphocyte detection mask on top of the same slide’s cancer-associated stroma mask could be used to calculate the number of TILs on a given slide. We could then divide this number by the area of cancer-associated stroma to find the associated density of TILs on the slide. By taking the “outer product” of cell and tissue types, we derived a wide array of composite features. In particular, we calculated counts, proportions, and densities of cells across different tissue types (e.g. density of macrophages in cancer-associated stroma versus in cancer tissue), under the hypothesis that these measures capture information that raw counts could not. To capture information regarding cell-cell proximity and interactions, we also calculated counts and proportions of each cell type within an 80 micron radius of each other cell type (e.g. count of lymphocytes within an 80 micron radius of fibroblasts). Cell-level counts, densities, and proportions comprised 264 HIFs. For each cell-tissue combination, we next applied the Birch clustering method (as implemented in the sklearn.cluster Python module) to partition cells into clusters^64^. To fit clustering structures as closely as possible to the spatial relationships found between cell types on the slide, we set a threshold of 100, a branching factor of 10, and allowed the algorithm to optimize the number of clusters returned. We used the returned clusters to calculate a series of features designed to capture spatial relationships between individual cells types within a given tissue type, including number of clusters, cluster size mean and standard deviation (SD), within-cluster dispersion mean and SD, cluster extent mean and SD, the Ball-Hall Index, and Calinski-Harabasz Index (n = 180 HIFs). For metrics where cluster exemplars were needed, the subcluster centers returned by the Birch algorithm were used.

### Patient-level aggregation

Patients with multiple tissue samples were represented by the single sample with the largest area of cancer tissue plus cancer-associated stroma, computed during tissue-based feature extraction. All subsequent analyses were conducted at the patient level.

### HIF clustering

Due to underlying biological relationships as well as the HIF generation process, there is significant correlation structure between many of the features. This presents a challenge of feature selection as much of the information contained in one feature will also be present in another. It also makes it difficult to control for multiple hypothesis testing, because the underlying number of tested hypotheses is significantly fewer than the number of features computed.

To identify groups of correlated HIFs, we clustered features via hierarchical agglomerative clustering using complete linkage, a cluster cutoff of 0.95, and pairwise correlation distance (1-absolute Spearman correlation) as the distance metric. We defined a set of HIF clusters for each cancer type independently, as well as another set for pan-cancer analyses (Supplemental Data 1). Clustering correlated features allows us to summarize the true underlying number of tested hypotheses.

### Visualization of cancer types in HIF space

Uniform Manifold Approximation and Projection (UMAP) was applied for dimensionality reduction and visualization of patient samples from the 607-dimension HIF space into two dimensions (using parameters: number of neighbors = 15, training epochs = 500, distance metric = euclidean). The V-Measure was computed to compare BRCA, STAD, SKCM, and NSCLC (LUAD and LUSC combined) classes against clusters generated by k-means (k = 4) applied to the 2-D UMAP projection^65,66^. To quantify differences between cancer types, HIF values were normalized pan-cancer into Z-scores. Median Z-scores were then computed per cancer type across twenty HIFs, each representing one of twenty HIF clusters defined pan-cancer. Representative HIFs were selected based on subjective interpretability and high variance across cancer types. To determine the statistical significance of median Z-scores that were greater in one cancer type relative to others, P-values were estimated with the one-sided Mann-Whitney U-test, considering NSCLC subtypes LUAD and LUSC as one type.

### Validation of HIFs against molecular markers

To validate the ability of HIFs to capture meaningful cell- and tissue-level information, we computed Spearman correlations between HIFs and four canonical immune markers from the PanImmune dataset^67^: (1) leukocyte infiltration, (2) IgG expression, (3) TGF-*β* expression, and (4) wound healing. Immune markers were quantified by mapping mRNA sequencing reads against gene sets associated with known immune expression signatures. To estimate the correlation between HIF clusters and immune markers, we computed the median absolute Spearman correlation per cluster and combined dependent P-values associated with individual correlations via the Empirical Brown’s method^35^. To control the false discovery rate, combined P-values per cluster were then corrected using the Benjamini-Hochberg procedure^68^. Correlation analyses were conducted for cancer types collectively and individually, using HIF clusters defined across all cancer types for assessment of concordance.

In addition, image-based cell quantifications for leukocyte fraction, lymphocyte fraction, and plasma cell fraction were validated by Spearman correlation to their sequencing-based equivalents from matched TCGA tumor samples, computed using CIBERSORT^67^. CIBERSORT (cell-type identification by estimating relative subsets of RNA transcripts) uses an immune signature matrix for deconvolution of observed RNA-Seq read counts into estimates of relative contributions between 22 immune cell profiles^34^.

### Molecular phenotype label curation

PD-1, PD-L1, and CTLA-4 expression data for each cancer type were collected from the PanImmune dataset^67^, while TIGIT expression data was collected from the National Cancer Institute Genomic Data Commons^69^. PD-1, PD-L1, CTLA-4, and TIGIT expression levels were quantified from mapped mRNA reads against genes PDCD1, CD274, CTLA-4, and TIGIT, respectively, and normalized as Z-scores across all cancer types in TCGA. Homologous recombination deficiency (HRD) scores were collected from Knijnenburg et al^70^. The HRD score was calculated as the sum of three components: 1) number of sub-chromosomal regions with allelic imbalance extending to the telomere, 2) number of chromosomal breaks between adjacent regions of least 10 Mb (mega base pairs), and 3) number of loss of heterozygosity regions of intermediate size (at least 15 Mb but less than whole chromosome length). Continuous immune checkpoint protein expression and HRD scores were binarized to high versus low classes using gaussian mixture model (GMM) clustering with unequal variance (Supplemental Figure 3). The binary threshold was defined as the intersection of the empirical densities between the two GMM-defined clusters. To evaluate the extent to which prediction tasks were correlated, Pearson correlation and percentage agreement metrics were computed pan-cancer (n = 1,893 patients) between the five molecular phenotypes in continuous and binarized form, respectively (Supplemental Figure 6).

### Hold-out set definition by TCGA tissue source site

TCGA provides tissue source site information, which denotes the medical institution or company that provided the patient sample. For each prediction task (described below), a hold-out set was defined as approximately 20-30% of patient samples obtained from sites not seen in the training set (Supplemental Table 4). This validation methodology enables us to demonstrate model generalizability across varying patient demographics and tissue collection processes intrinsic to different tissue source sites. Patient barcodes corresponding to hold-out and training sets are provided in Supplemental Data 2.

### Supervised prediction of molecular phenotypes

We conducted supervised prediction of binarized high versus low expression of five clinically-relevant phenotypes: (1) PD-1 expression, (2) PD-L1 expression, (3) CTLA-4 expression, (4) HRD score, and (5) TIGIT expression. Predictions were conducted pan-cancer as well as for cancer types individually. SKCM was excluded from prediction tasks 1-4 due to insufficient outcome labels (number of observations < 100 for tasks 1-3; number of positive labels < 10 for task 4). For each prediction task, we trained a logistic sparse group lasso (SGL) model71 tuned by nested cross validation (CV) with three outer folds and five inner folds using the corresponding training set. SGL provides regularization at both an individual covariate (as in traditional lasso) and user-defined group level, thus encouraging group-wise and within group sparsity. The HIF clusters defined per cancer type and pancancer (previously described) were inputted as groups. HIFs were normalized to mean = 0 and SD = 1. In accordance with nested CV, hyper-parameter tuning was conducted using the inner loops and mean generalization error and variance were estimated from the outer loops. The three tuned models, each trained on two of the three outer folds and evaluated on the third outer fold, were ensembled by averaging predicted probabilities for final evaluation (reported in Figure 6a; Supplemental Table 5) on the hold-out set. Holdout performance was evaluated by AUROC and AUPRC. To identify predictive features, beta values from the three outer fold models were averaged to obtain ensemble beta values per HIF (see Figure 6b caption for more details).

### Statistical analysis

To compute 95% confidence intervals for each prediction task, we generated empirical distributions of AUROC and AUPRC metrics each consisting of 1000 bootstrapped metrics. Bootstrapped metrics were obtained by sampling with replacement from matched model predictions (probabilities) and true labels for the corresponding hold-out set, and re-computing AUROC and AUPRC on these two bootstrapped vectors. P-values for ensemble beta values of predictive HIFs were computed using a permutation test with 1000 iterations. During each iteration, labels in the training set were permuted and the previously described training process of nested CV and ensembling was re-applied to generate a new set of ensemble beta values per HIF. P-values for individual HIFs were then obtained by comparing beta values in the original ensemble model against the corresponding null distribution of ensemble beta values. Individual HIF P-values were combined into cluster-level P-values via the Empirical Brown’s method35 and corrected using the Benjamini-Hochberg procedure^68^.

### Data availability

The Cancer Genome Atlas dataset may be accessed at https://www.cancer.gov/about-nci/organization/ccg/research/structural-genomics/tcga. The relevant data consists of 2,917 hematoxylin and eosin-stained WSIs of breast cancer, non-small cell lung adenocarcinoma, non-small cell lung squamous cell carcinoma, gastric adenocarcinoma, and skin cutaneous melanoma specimens from 2,634 patients.

### Code availability

The source code used to generate figures in this work can be downloaded from: https://github.com/Path-AI/hif2gene.

