## Supplemental Information for "Dense, high-resolution mapping of cells and tissues from pathology images for the interpretable prediction of molecular phenotypes in cancer"

**Supplemental Table 1. Patient demographics and characteristics**

Whole-slide images (WSIs) and clinical data were acquired from The Cancer Genome Atlas (TCGA) and encompassed BRCA, LUAD, LUSC, SKCM, and STAD patients (n=2634 total patients) from 95 distinct clinical sites. Patients with multiple samples were represented by the sample with the largest tumor region.

| | SKCM | | STAD | | BRCA | | LUSC | | LUAD | |
|---|---|---|---|---|---|---|---|---|---|---|
| **Number of Patients** | 327 | | 407 | | 1044 | | 412 | | 444 | |
| **Number of Distinct Tissue Source Sites** | 24 | | 22 | | 39 | | 34 | | 32 | |
| **Age at Diagnosis (Mean +/- SD)** | 57.4 +/- 15.8 | | 64.8 +/- 10.6 | | 58.5 +/- 13.2 | | 67.7 +/- 8.6 | | 65.1 +/- 10.1 | |
| **Gender (%)** | | | | | | | | | | |
| Male | 200 | 61.16% | 269 | 66.09% | 12 | 1.15% | 306 | 74.27% | 205 | 46.17% |
| Female | 127 | 38.84% | 138 | 33.91% | 1032 | 98.85% | 106 | 25.73% | 239 | 53.83% |
| **Stage (%)** | | | | | | | | | | |
| 0 | 4 | 1.22% | 0 | 0.00% | 0 | 0.00% | 0 | 0.00% | 0 | 0.00% |
| 1 | 55 | 16.82% | 92 | 22.60% | 188 | 18.01% | 214 | 51.94% | 265 | 59.68% |
| 2 | 83 | 25.38% | 122 | 29.98% | 595 | 56.99% | 126 | 30.58% | 107 | 24.10% |
| 3 | 129 | 39.45% | 183 | 44.96% | 238 | 22.80% | 68 | 16.50% | 65 | 14.64% |
| 4 | 14 | 4.28% | 0 | 0.00% | 0 | 0.00% | 0 | 0.00% | 0 | 0.00% |
| 1-2 Not Otherwise Specified | 12 | 3.67% | 0 | 0.00% | 0 | 0.00% | 0 | 0.00% | 0 | 0.00% |
| Unknown or Discrepancy | 30 | 9.17% | 10 | 2.46% | 23 | 2.20% | 4 | 0.97% | 7 | 1.58% |

**Supplemental Table 2. Cell- and tissue-type annotation counts**
Annotations for six cell classes and four tissue classes were obtained from a multi-hundred pathologist network. Cell type annotations are individual pixels while tissue region annotations are fully connected components of pixels. The total number of cell and tissue type annotations collected on TCGA H&E images and several additional H&E datasets in aggregate are shown broken down by cancer type and annotation type. In total, 5,719 WSIs curated from TCGA and additional datasets were used during development of the deep learning models (training and validation). Tissue-level segmentations and cell-level predictions were then generated on 2,826 TCGA WSIs.

| **TCGA Datasets** | | | | | | |
|---|---|---|---|---|---|---|
| **Cell-Level Annotations** | BRCA | LUAD | LUSC | SKCM | STAD | Total |
| Cancer Cell | 25561 | 5077 | 24117 | 43200 | 18949 | 116904 |
| Fibroblast | 7464 | 3367 | 8833 | 10994 | 9155 | 39813 |
| Lymphocyte | 7904 | 4223 | 9211 | 17247 | 11388 | 49973 |
| Macrophage | 3062 | 3474 | 7758 | 5596 | 4297 | 24187 |
| Plasma Cell | 1792 | 2689 | 6997 | 5990 | 5183 | 22651 |
| Other Cell / Background | 12057 | 3128 | 11632 | 22069 | 20626 | 69512 |
| Total | 57840 | 21958 | 68548 | 105096 | 69598 | **323040** |
| **Tissue-Level Annotations** | | | | | | |
| Cancer Tissue | 3660 | 2549 | 5433 | 7797 | 3800 | 23239 |
| Cancer-Associated Stroma | 2001 | 1696 | 4591 | 2287 | 2170 | 12745 |
| Necrosis | 152 | 318 | 1335 | 991 | 431 | 3227 |
| Normal / Background | 1824 | 992 | 2467 | 8536 | 2516 | 16335 |
| Total | 7637 | 5555 | 13826 | 19611 | 8917 | **55546** |
| **Whole-Slide Images** | | | | | | |
| Development | 359 | 137 | 307 | 358 | 400 | 1561 |
| Inference | 1117 | 501 | 443 | 358 | 407 | 2826 |
| QC Excluded | 20 | 37 | 24 | 5 | 5 | 91 |
| Total | 1137 | 538 | 467 | 363 | 412 | **2917** |
| **Additional Datasets** | | | | | | |
| **Cell-Level Annotations** | BRCA | LUAD | LUSC | SKCM | STAD | Total |
| Cancer Cell | 40518 | 101979 | 0 | 254392 | 9435 | 406324 |
| Fibroblast | 9201 | 38603 | 2773 | 49815 | 3106 | 103498 |
| Lymphocyte | 23145 | 65223 | 21852 | 125842 | 3816 | 239878 |
| Macrophage | 5422 | 17424 | 9723 | 49879 | 987 | 83435 |

| | | | | | | |
|---|---|---|---|---|---|---|
| Plasma Cell | 6797 | 6401 | 5487 | 19158 | 1091 | 38934 |
| Other Cell / Background | 45391 | 28336 | 5226 | 159118 | 9564 | 247635 |
| Total | 130474 | 257966 | 45061 | 658204 | 27999 | **1119704** |
| **Tissue-Level Annotations** | | | | | | |
| Cancer Tissue | 9945 | 14654 | 6471 | 27889 | 4428 | 63387 |
| Cancer-Associated Stroma | 6617 | 8826 | 3866 | 14076 | 2257 | 35642 |
| Necrosis | 741 | 1279 | 1256 | 2843 | 326 | 6445 |
| Normal / Background | 5791 | 14436 | 2752 | 16446 | 1119 | 40544 |
| Total | 23094 | 39195 | 14345 | 61254 | 8130 | **146018** |
| **Whole-Slide Images** | | | | | | |
| Development | 698 | 1908 | 438 | 1002 | 112 | 4158 |
| Inference | 0 | 0 | 0 | 0 | 0 | 0 |
| Total | 698 | 1908 | 438 | 1002 | 112 | **4158** |
| **Combined** | | | | | | |
| | BRCA | LUAD | LUSC | SKCM | STAD | Total |
| **Total Cell-Level Annotations** | 188314 | 279924 | 113609 | 763300 | 97597 | **1442744** |
| **Total Tissue-Level Annotations** | 30731 | 44750 | 28171 | 80865 | 17047 | **201564** |
| **Total Whole-Slide Images** | 1835 | 2446 | 905 | 1365 | 524 | **7075** |

**Supplemental Table 3. Validation of human-interpretable image features (HIFs) against immune markers broken down by cancer type**

The top three HIF clusters, ranked by median absolute Spearman correlation ρ (computed across HIFs in a given cluster), for each of four immune markers (leukocyte infiltration, IgG expression, TGF-β expression, wound healing signature) and a negative control (Case ID). Interquartile ranges (IQR) are also shown. Q-values are combined (via the Empirical Brown's method) and corrected (via Benjamini-Hochberg procedure) P-values per cluster. SKCM-specific analyses were excluded due to insufficient sample sizes. Pan-cancer analyses, however, include SKCM patients. Clusters were defined pan-cancer to enable comparisons across cancer types. Notably, we can observe high concordance among the top-correlated HIF clusters per immune marker across cancer types. HIF clusters are provided in Supplemental Data 1.

| **Leukocyte Infiltration Signature** | | | |
|---|---|---|---|
| Cancer Type | HIF Cluster | Median (IQR) Abs(ρ) | Q-Value |
| BRCA (n=1022) | 10 | 0.48 (0.45, 0.51) | 1.37E-125 |
| | 9 | 0.41 (0.36, 0.47) | 3.85E-166 |
| | 7 | 0.37 (0.33, 0.43) | 8.97E-141 |
| LUAD (n=390) | 10 | 0.49 (0.41, 0.51) | 5.00E-23 |
| | 9 | 0.39 (0.29, 0.43) | 1.07E-12 |
| | 7 | 0.27 (0.17, 0.33) | 1.86E-29 |
| LUSC (n=400) | 9 | 0.40 (0.36, 0.44) | 6.79E-13 |
| | 10 | 0.38 (0.33, 0.43) | 4.06E-12 |
| | 4 | 0.22 (0.20, 0.27) | 3.51E-17 |
| STAD (n=328) | 10 | 0.36 (0.32, 0.43) | 1.27E-11 |
| | 8 | 0.35 (0.32, 0.38) | 6.42E-08 |
| | 12 | 0.27 (0.23, 0.33) | 1.31E-07 |
| Pan-Cancer (n=2202) | 10 | 0.48 (0.45, 0.53) | 8.48E-279 |
| | 9 | 0.46 (0.43, 0.49) | 2.09E-280 |
| | 7 | 0.40 (0.33, 0.42) | 1.09E-295 |

| **IgG Expression** | | | |
|---|---|---|---|
| Cancer Type | HIF Cluster | Median (IQR) Abs(ρ) | Q-Value |

| | | | |
|---|---|---|---|
| BRCA (n=1022) | 9 | 0.44 (0.40, 0.49) | 3.05E-194 |
| | 10 | 0.44 (0.41, 0.46) | 6.62E-100 |
| | 8 | 0.37 (0.20, 0.41) | 3.58E-171 |
| LUAD (n=390) | 9 | 0.34 (0.23, 0.40) | 1.30E-09 |
| | 10 | 0.25 (0.22, 0.27) | 2.14E-08 |
| | 4 | 0.17 (0.13, 0.20) | 7.04E-10 |
| LUSC (n=400) | 9 | 0.36 (0.29, 0.44) | 1.03E-15 |
| | 10 | 0.26 (0.18, 0.31) | 5.09E-09 |
| | 4 | 0.24 (0.18, 0.26) | 1.03E-15 |
| STAD (n=328) | 8 | 0.35 (0.32, 0.40) | 2.39E-09 |
| | 9 | 0.32 (0.24, 0.37) | 1.14E-06 |
| | 10 | 0.26 (0.23, 0.29) | 7.62E-05 |
| Pan-Cancer (n=2202) | 9 | 0.58 (0.53, 0.60) | 2.67E-204 |
| | 7 | 0.44 (0.38, 0.49) | 4.83E-54 |
| | 10 | 0.44 (0.38, 0.47) | 4.86E-42 |

| **TGF-β Signature** | | | |
|---|---|---|---|
| Cancer Type | HIF Cluster | Median (IQR) Abs(ρ) | Q-Value |
| BRCA (n=1022) | 12 | 0.37 (0.24, 0.38) | 7.30E-85 |
| | 14 | 0.35 (0.27, 0.43) | 3.90E-75 |
| | 4 | 0.32 (0.22, 0.35) | 1.55E-195 |
| LUAD (n=390) | 4 | 0.26 (0.22, 0.28) | 5.57E-10 |
| | 14 | 0.23 (0.10, 0.25) | 1.38E-06 |
| | 12 | 0.20 (0.16, 0.23) | 3.85E-06 |
| LUSC (n=400) | 14 | 0.30 (0.26, 0.36) | 1.81E-10 |
| | 12 | 0.27 (0.23, 0.30) | 4.66E-13 |
| | 4 | 0.25 (0.16, 0.26) | 2.04E-08 |
| STAD (n=328) | 12 | 0.40 (0.36, 0.41) | 7.00E-16 |
| | 14 | 0.33 (0.26, 0.38) | 3.26E-06 |
| | 4 | 0.31 (0.21, 0.36) | 5.60E-12 |
| Pan-Cancer (n=2202) | 4 | 0.31 (0.21, 0.32) | 1.03E-90 |

| | | | |
|---|---|---|---|
| | 14 | 0.28 (0.24, 0.31) | 4.09E-104 |
| | 12 | 0.25 (0.22, 0.29) | 7.62E-104 |

| **Wound Healing Signature** | | | |
|---|---|---|---|
| Cancer Type | HIF Cluster | Median (IQR) Abs(ρ) | Q-Value |
| BRCA (n=1022) | 16 | 0.37 (0.36, 0.42) | 9.84E-76 |
| | 2 | 0.25 (0.29, 0.41) | 3.42E-74 |
| | 9 | 0.29 (0.16, 0.35) | 3.42E-74 |
| LUAD (n=390) | 16 | 0.15 (0.14, 0.18) | 0.19 |
| | 2 | 0.13 (0.08, 0.19) | 9.39E-08 |
| | 20 | 0.10 (0.07, 0.14) | 1.84E-02 |
| LUSC (n=400) | 2 | 0.19 (0.14, 0.22) | 1.56E-04 |
| | 12 | 0.11 (0.10, 0.15) | 1.87E-03 |
| | 1 | 0.09 (0.07, 0.11) | 1.82E-02 |
| STAD (n=328) | 12 | 0.42 (0.41, 0.47) | 1.73E-18 |
| | 4 | 0.40 (0.24, 0.47) | 4.76E-20 |
| | 14 | 0.32 (0.28, 0.38) | 6.29E-08 |
| Pan-Cancer (n=2202) | 16 | 0.35 (0.27, 0.38) | 1.80E-167 |
| | 2 | 0.34 (0.25, 0.40) | 3.85E-141 |
| | 12 | 0.29 (0.24, 0.36) | 7.59E-96 |

| **Case ID (Negative Control)** | | | |
|---|---|---|---|
| Cancer Type | HIF Cluster | Median (IQR) Abs(ρ) | Q-Value |
| BRCA (n=1022) | 17 | 0.07 (0.06, 0.07) | 0.36 |
| | 19 | 0.05 (0.04, 0.06) | 0.68 |
| | 13 | 0.04 (0.02, 0.04) | 0.86 |
| LUAD (n=390) | 19 | 0.17 (0.10, 0.21) | 0.20 |
| | 7 | 0.06 (0.03, 0.08) | 0.78 |
| | 14 | 0.05 (0.03, 0.07) | 0.78 |
| LUSC (n=400) | 13 | 0.10 (0.01, 0.12) | 0.62 |
| | 1 | 0.09 (0.07, 0.11) | 0.26 |

| | | | |
|---|---|---|---|
| | 2 | 0.08 (0.07, 0.14) | 0.88 |
| STAD (n=328) | 18 | 0.07 (0.04, 0.07) | 0.92 |
| | 1 | 0.07 (0.04, 0.09 | 0.99 |
| | 12 | 0.06 (0.03, 0.08) | 0.92 |
| Pan-Cancer (n=2202) | 17 | 0.06 (0.04, 0.06) | 0.08 |
| | 19 | 0.05 (0.03, 0.05) | 0.08 |
| | 5 | 0.02 (0.01, 0.04) | 0.40 |

**Supplemental Table 4. Holdout set definition**

Hold-out sets were defined as approximately 20-30% of all samples and consisted of 2-3 tissue source sites not included in the training set. This enabled us to evaluate our final ensemble models on varying patient demographics and institutions. The pan-cancer hold-out set was defined as the concatenation of hold-out sets from individual cancer types. The percentages of positive labels in the training and hold-out sets are shown. For a given cancer type, the same set of hold-out tissue source sites was used across prediction outcomes.

| Cancer Type | Tissue Source Site / Code | Hold-Out N | Immune Score | % Positive Train | % Positive Hold-Out |
|---|---|---|---|---|---|
| BRCA | BH (University of Pittsburgh)<br>A2 (Walter Reed) | 239/1022 (23.4%) | PD-1 | 45.8% | 48.1% |
| | | | PDL-1 | 47.9% | 51.0% |
| | | | CTLA-4 | 53.3% | 51.5% |
| | | 208/904 (23.0%) | HRD | 16.7% | 12.5% |
| | | 201/977 (20.6%) | TIGIT | 48.8% | 49.3% |
| LUAD | 55 (International Genomics Consortium)<br>50 (University of Pittsburgh) | 95/390 (24.4%) | PD-1 | 46.1% | 51.6% |
| | | | PDL-1 | 57.6% | 63.2% |
| | | | CTLA-4 | 51.9% | 53.7% |
| | | 104/417 (24.9%) | HRD | 26.2% | 19.2% |
| | | 100/420 (23.8%) | TIGIT | 50.3% | 60.0% |
| LUSC | 85 (Asterand)<br>66 (Indivumed)<br>33 (Johns Hopkins) | 98/400 (24.5%) | PD-1 | 48.7% | 42.9% |
| | | | PDL-1 | 35.8% | 36.7% |
| | | | CTLA-4 | 34.7% | 37.7% |
| | | 96/384 (25.0%) | HRD | 18.8% | 19.8% |
| | | 98/398 (24.6%) | TIGIT | 33.0% | 30.6% |
| SKCM | D3 (MD Anderson)<br>WE (Norfolk and Norwich Hospital) | 83/316 (26.3%) | TIGIT | 57.5% | 63.9% |
| STAD | VQ (Barretos Cancer Hospital)<br>RD (Peter MacCallum Cancer Center)<br>FP (International Genomics Consortium) | 87/328 (26.5%) | PD-1 | 52.7% | 49.4% |
| | | | PDL-1 | 42.3% | 42.5% |
| | | | CTLA-4 | 61.4% | 65.5% |
| | | 80/316 (25.3%) | HRD | 28.0% | 50.0% |
| | | 87/341 (25.5%) | TIGIT | 66.9% | 63.2% |
| Pan-Cancer | Concatenated (BRCA, LUAD, LUSC, STAD) | 519/2140 (24.3%) | PD-1 | 47.4% | 48.0% |
| | | | PDL-1 | 46.6% | 49.1% |
| | | | CTLA-4 | 51.3% | 51.1% |
| | | 488/2021 (24.1%) | HRD | 20.7% | 21.5% |
| | Concatenated (BRCA, LUAD, LUSC, SKCM, STAD) | 569/2452 (23.2%) | TIGIT | 50.1% | 52.2% |

**Supplemental Table 5. Full cross-validation and hold-out results from predicting clinically-relevant phenotypes using HIFs**

Supervised prediction was conducted on binarized high versus low expression of five clinically-relevant molecular phenotypes: (1) PD-1 expression, (2) PD-L1 expression, (3) CTLA-4 expression, (4) HRD score, and (5) TIGIT expression. SKCM predictions were conducted only for TIGIT expression due to insufficient labels for the remaining phenotypes. Pan-cancer analyses used the same binary labels thresholded independently by cancer type. Pan-cancer predictions for TIGIT included all five cancer types, while pan-cancer predictions for the remaining phenotypes included the four cancer types excluding SKCM. Cross-validation (CV) and hold-out performance were measured by area under the receiver operating characteristic (AUROC) and area under the precision-recall curve (AUPRC). Random classifiers correspond to AUROC=0.50 and AUPRC=percentage of positive labels. CV metrics are presented as the mean and standard deviation computed across three models, each trained on two of the three outer folds and evaluated on the third in accordance with nested CV. Empirical bootstrapping using 1000 resamples was applied to compute 95% confidence intervals for hold-out metrics. Sample sizes for training and hold-out sets are shown. Notably, immune checkpoint protein, HRD, and TIGIT labels were curated from distinct datasets, resulting in differing sample size numbers.

| | | Cross Validation Metrics | | | Hold-Out Metrics | | |
|---|---|---|---|---|---|---|---|
| Cancer Type (Training, Hold-Out) | Immune Score | Mean AUROC | Mean AUPRC | % Positive | AUROC (95% CI) | AUPRC (95% CI) | % Positive |
| BRCA n=783, 239 | PD-1 | 0.773 +/- 0.01 | 0.757 +/- 0.01 | 45.8% | 0.777 (0.716, 0.835) | 0.778 (0.696, 0.851) | 48.1% |
| | PD-L1 | 0.729 +/- 0.02 | 0.714 +/- 0.01 | 47.9% | 0.663 (0.594, 0.733) | 0.702 (0.618, 0.775) | 51.0% |
| | CTLA-4 | 0.842 +/- 3E-3 | 0.860 +/- 5E-3 | 53.3% | 0.794 (0.739, 0.848) | 0.818 (0.752, 0.875) | 51.5% |
| n=696, 208 | HRD | 0.743 +/- 0.01 | 0.393 +/- 0.02 | 16.7% | 0.774 (0.665, 0.867) | 0.354 (0.196, 0.570) | 12.5% |
| n=776, 201 | TIGIT | 0.793 +/- 0.02 | 0.804 +/- 0.01 | 48.8% | 0.788 (0.723, 0.848) | 0.799 (0.724, 0.865) | 49.3% |
| LUAD n=295, 95 | PD-1 | 0.814 +/- 0.03 | 0.786 +/- 0.02 | 46.1% | 0.705 (0.593, 0.805) | 0.726 (0.587, 0.849) | 51.6% |
| | PD-L1 | 0.811 +/- 0.03 | 0.849 +/- 0.02 | 57.6% | 0.727 (0.610, 0.832) | 0.776 (0.653, 0.903) | 63.2% |
| | CTLA-4 | 0.781 +/- 0.02 | 0.802 +/- 0.01 | 51.9% | 0.711 (0.605, 0.808) | 0.756 (0.634, 0.849) | 53.7% |
| n=313 ,104 | HRD | 0.594 +/- 0.04 | 0.325 +/- 0.06 | 26.2% | 0.549 (0.406, 0.685) | 0.209 (0.119, 0.375) | 19.2% |
| n= 320, 100 | TIGIT | 0.732 +/- 0.03 | 0.734 +/- 0.05 | 50.3% | 0.766 (0.670, 0.852) | 0.848 (0.766, 0.911) | 60.0% |
| LUSC n=302, 98 | PD-1 | 0.766 +/- 0.04 | 0.773 +/- 0.04 | 48.7% | 0.675 (0.571, 0.775) | 0.611 (0.459, 0.745) | 42.9% |
| | PD-L1 | 0.650 +/- 0.02 | 0.518 +/- 0.02 | 35.8% | 0.513 (0.394, 0.637) | 0.425 (0.292, 0.575) | 36.7% |
| | CTLA-4 | 0.760 +/- 0.01 | 0.663 +/- 0.01 | 34.7% | 0.689 | 0.541 | 37.7% |

| | | | | | | | |
|---|---|---|---|---|---|---|---|
| | | | | | (0.571, 0.798) | (0.377, 0.706) | |
| n=288, 96 | HRD | 0.564 +/- 0.02 | 0.228 +/- 0.05 | 18.8% | 0.476 (0.338, 0.606) | 0.178 (0.106, 0.295) | 19.8% |
| n=300, 98 | TIGIT | 0.703 +/- 0.01 | 0.559 +/- 0.03 | 33.0% | 0.706 (0.597, 0.811) | 0.494 (0.329, 0.679) | 30.6% |
| SKCM n=233, 83 | TIGIT | 0.854 +/- 0.05 | 0.895 +/- 0.03 | 57.5% | 0.864 (0.771, 0.954) | 0.871 (0.757, 0.976) | 63.9% |
| STAD n=241, 87 | PD-1 | 0.826 +/- 0.02 | 0.847 +/- 0.01 | 52.7% | 0.684 (0.566, 0.790) | 0.701 (0.541, 0.825) | 49.4% |
| | PD-L1 | 0.838 +/- 0.06 | 0.794 +/- 0.06 | 42.3% | 0.601 (0.470, 0.733) | 0.616 (0.457, 0.767) | 42.5% |
| | CTLA-4 | 0.799 +/- 0.03 | 0.856 +/- 0.01 | 61.4% | 0.626 (0.498, 0.746) | 0.759 (0.638, 0.867) | 65.5% |
| n=236, 80 | HRD | 0.681 +/- 0.03 | 0.433 +/- 0.07 | 28.0% | 0.628 (0.501, 0.750) | 0.647 (0.494, 0.797) | 50.0% |
| n=254, 87 | TIGIT | 0.674 +/- 0.03 | 0.764 +/- 0.03 | 66.9% | 0.657 (0.536, 0.779) | 0.756 (0.617, 0.882) | 63.2% |
| Pan-Cancer n=1621, 519 | PD-1 | 0.718 +/- 0.03 | 0.691 +/- 0.04 | 47.4% | 0.677 (0.636, 0.723) | 0.673 (0.615, 0.731) | 48.0% |
| | PD-L1 | 0.670 +/- 0.04 | 0.643 +/- 0.04 | 46.6% | 0.626 (0.574, 0.674) | 0.639 (0.574, 0.707) | 49.1% |
| | CTLA-4 | 0.727 +/- 0.05 | 0.732 +/- 0.06 | 51.3% | 0.738 (0.697, 0.785) | 0.752 (0.699, 0.807) | 51.1% |
| n=1533, 488 | HRD | 0.640 +/- 0.02 | 0.293 +/- 0.02 | 20.7% | 0.651 (0.596, 0,709) | 0.336 (0.263, 0.421) | 21.5% |
| n=1883, 569 | TIGIT | 0.771 +/- 0.01 | 0.770 +/- 0.01 | 50.1% | 0.792 (0.754, 0.827) | 0.799 (0.750, 0.844) | 52.2% |

**Supplemental Table 6. Combined and corrected P-values for predictive HIF clusters**

P-values associated with individual HIFs were computed based on a 1000-iteration permutation test. Individual HIF P-values were combined via the Empirical Brown's Method into cluster-level P-values and corrected using the Benjamini-Hochberg procedure (see Statistical Analysis in Methods for more details). Q-values (combined and corrected cluster-level P-values) are shown broken down by cancer type and prediction task. HIF clusters with no non-zero ensemble betas are grayed. The top five predictive HIF clusters per prediction (or fewer in the case of ensemble models with <5 non-zero clusters) are bolded.

| Pan-Cancer | | | | | |
|---|---|---|---|---|---|
| Cluster Number | PD-1 Prediction Q-Value | PD-L1 Prediction Q-Value | CTLA-4 Prediction Q-Value | HRD Prediction Q-Value | TIGIT Prediction Q-Value |
| 1 | **1.64E-03** | **3.51E-02** | **5.00E-02** | | **5.84E-02** |
| 2 | | **2.64E-04** | 4.90E-05 | **2.26E-04** | |
| 3 | 2.90E-01 | 3.80E-03 | 2.16E-04 | | **1.38E-03** |
| 4 | | 2.63E-02 | 1.72E-06 | **4.36E-12** | |
| 5 | | **7.04E-08** | 8.17E-05 | | |
| 6 | 1.50E-02 | 7.62E-02 | 1.78E-11 | | |
| 7 | **2.40E-01** | **4.84E-03** | 2.11E-01 | | |
| 8 | **1.20E-01** | 7.84E-01 | 6.22E-03 | | |
| 9 | **1.40E-02** | 3.74E-01 | 4.87E-01 | **6.94E-04** | |
| 10 | **1.24E-03** | **5.14E-04** | **2.65E-04** | | **6.49E-02** |
| 11 | | 8.36E-03 | **3.80E-07** | | |
| 12 | | 4.79E-03 | 2.67E-04 | **3.52E-12** | |
| 13 | | 9.41E-08 | 1.58E-10 | 1.82E-06 | |
| 14 | | 5.95E-04 | 7.22E-05 | | |
| 15 | | | **5.27E-20** | | |
| 16 | 5.88E-01 | 6.12E-01 | 1.15E-06 | **1.88E-04** | |
| 17 | | | | | |
| 18 | | | | | |
| 19 | | 1.71E-02 | 7.17E-07 | | |
| 20 | | 4.84E-03 | **3.18E-05** | | |

| BRCA |
|---|

| Cluster Number | PD-1 Prediction Q-Value | PD-L1 Prediction Q-Value | CTLA-4 Prediction Q-Value | HRD Prediction Q-Value | TIGIT Prediction Q-Value |
|---|---|---|---|---|---|
| 1 | **9.51E-02** | **4.42E-02** | 9.05E-03 | 1.55E-02 | **3.15E-01** |
| 2 | | 7.49E-03 | 9.05E-03 | **4.12E-04** | |
| 3 | | 1.28E-05 | **4.46E-10** | 1.65E-07 | 1.18E-03 |
| 4 | | 1.12E-01 | 6.67E-08 | 3.08E-05 | **1.34E-01** |
| 5 | | 6.90E-02 | 1.73E-02 | 1.65E-02 | 7.30E-01 |
| 6 | | 5.10E-04 | | 9.06E-07 | |
| 7 | | 1.02E-08 | 2.24E-03 | 1.91E-03 | **1.98E-03** |
| 8 | **9.51E-02** | **4.13E-07** | **1.72E-05** | 3.43E-05 | **7.53E-05** |
| 9 | **1.60E-05** | **1.10E-04** | **2.47E-05** | 1.50E-03 | **1.59E-02** |
| 10 | | 1.86E-09 | **7.54E-13** | 4.32E-02 | 2.10E-07 |
| 11 | | 6.60E-03 | 9.05E-03 | **1.50E-03** | 4.69E-01 |
| 12 | | | 3.21E-04 | 1.56E-02 | 3.15E-01 |
| 13 | | **3.03E-07** | 3.35E-06 | **7.37E-06** | 4.53E-01 |
| 14 | | 3.54E-03 | **2.12E-16** | **2.32E-13** | 6.87E-05 |
| 15 | | 5.10E-04 | 1.20E-02 | 1.10E-04 | 1.49E-01 |
| 16 | | 2.66E-02 | 8.25E-08 | 6.64E-05 | 9.77E-01 |
| 17 | | | | | |
| 18 | | 8.23E-02 | 4.87E-02 | | 1.53E-01 |
| 19 | | **1.89E-03** | | | |
| 20 | | | 2.05E-04 | **3.43E-04** | 3.15E-01 |

| LUAD | | | | |
|---|---|---|---|---|
| Cluster Number | PD-1 Prediction Q-Value | PD-L1 Prediction Q-Value | CTLA-4 Prediction Q-Value | TIGIT Prediction Q-Value |
| 1 | 6.62E-03 | 1.86E-04 | 4.08E-04 | **1.62E-03** |
| 2 | | | | **1.05E-04** |
| 3 | 3.67E-02 | **6.50E-03** | 1.88E-03 | 2.45E-01 |
| 4 | **4.77E-04** | 4.58E-02 | 1.07E-04 | 2.79E-04 |
| 5 | 1.72E-03 | 1.80E-01 | 3.25E-03 | 4.53E-01 |
| 6 | **7.44E-09** | 2.87E-01 | 1.99E-03 | 3.46E-01 |
| 7 | 2.06E-01 | 3.69E-02 | 6.63E-02 | 4.53E-01 |
| 8 | 5.73E-02 | | **3.00E-06** | **1.44E-01** |
| 9 | 1.46E-03 | | | 1.84E-02 |

| | | | | |
|---|---|---|---|---|
| 10 | 9.91E-02 | 1.08E-01 | 1.96E-02 | 4.67E-01 |
| 11 | 2.59E-02 | 2.33E-03 | 6.01E-01 | 2.71E-01 |
| 12 | 5.73E-02 | 4.00E-03 | 4.89E-04 | 3.62E-01 |
| 13 | **3.05E-05** | **1.86E-04** | **3.00E-06** | **1.90E-03** |
| 14 | 8.88E-01 | 7.25E-02 | 2.19E-02 | 1.03E-01 |
| 15 | 2.79E-01 | **4.44E-05** | **3.00E-06** | 3.66E-01 |
| 16 | | | | 1.07E-03 |
| 17 | | | 1.74E-01 | 1.09E-02 |
| 18 | **1.51E-04** | | **7.95E-03** | 4.78E-01 |
| 19 | **2.43E-03** | **2.33E-03** | **1.89E-03** | **6.67E-03** |
| 20 | 1.60E-01 | **4.23E-08** | 5.00E-05 | 5.57E-02 |

| LUSC | | | | SKCM | |
|---|---|---|---|---|---|
| Cluster Number | PD-1 Prediction Q-Value | CTLA-4 Prediction Q-Value | TIGIT Prediction Q-Value | Cluster Number | TIGIT Prediction Q-Value |
| 1 | 1.30E-01 | | 3.51E-01 | 1 | |
| 2 | 1.89E-03 | | | 2 | **2.70E-10** |
| 3 | **2.51E-06** | 2.88E-01 | 1.80E-02 | 3 | 7.87E-02 |
| 4 | 1.16E-02 | | 2.74E-01 | 4 | 1.45E-02 |
| 5 | 4.42E-03 | | 6.70E-01 | 5 | |
| 6 | 3.30E-02 | | 5.42E-01 | 6 | |
| 7 | 1.89E-03 | | 4.00E-01 | 7 | |
| 8 | 1.95E-02 | **1.87E-02** | 2.11E-02 | 8 | **1.45E-02** |
| 9 | 7.18E-04 | 9.13E-01 | **5.61E-03** | 9 | |
| 10 | 2.03E-02 | | 5.41E-01 | 10 | 2.83E-01 |
| 11 | 4.09E-03 | 2.88E-01 | 4.66E-02 | 11 | |
| 12 | 1.21E-02 | | 4.02E-01 | 12 | |
| 13 | 1.89E-03 | | 3.77E-02 | 13 | |
| 14 | **1.46E-02** | **1.49E-01** | **3.73E-02** | 14 | 8.18E-02 |
| 15 | 2.44E-02 | | | 15 | |
| 16 | 3.12E-02 | | 2.75E-01 | 16 | |
| 17 | 3.28E-02 | | 6.70E-01 | 17 | |
| 18 | **2.62E-08** | | **1.36E-03** | 18 | |
| 19 | 2.66E-02 | | | 19 | **1.28E-06** |
| 20 | 1.32E-01 | | **5.61E-03** | 20 | **2.68E-04** |

| | | | | | |
|---|---|---|---|---|---|
| 21 | **2.56E-03** | **8.00E-03** | **1.36E-03** | 21 | |
| 22 | 5.96E-02 | | | 22 | **5.19E-04** |
| 23 | **1.33E-04** | | | 23 | |
| 24 | 9.96E-02 | **1.33E-02** | 8.27E-03 | 24 | |
| 25 | 1.07E-01 | **8.00E-03** | | | |

| STAD | | | | | |
|---|---|---|---|---|---|
| Cluster Number | PD-1 Prediction Q-Value | PD-L1 Prediction Q-Value | CTLA-4 Prediction Q-Value | HRD Prediction Q-Value | TIGIT Prediction Q-Value |
| 1 | 1.47E-01 | 2.65E-05 | **2.13E-03** | 5.93E-02 | 1.28E-03 |
| 2 | 7.90E-01 | **7.26E-06** | 9.76E-02 | **1.50E-05** | 4.69E-04 |
| 3 | 1.47E-01 | 4.89E-04 | 6.43E-03 | | 8.40E-04 |
| 4 | 7.51E-01 | 2.40E-04 | 1.24E-02 | | 4.40E-02 |
| 5 | | 7.60E-08 | 1.86E-02 | **1.50E-05** | 3.24E-04 |
| 6 | | 1.88E-07 | 6.20E-07 | 5.85E-02 | 3.24E-04 |
| 7 | | **7.19E-05** | 7.16E-01 | | 5.76E-04 |
| 8 | | 5.95E-06 | 2.49E-01 | | 2.18E-03 |
| 9 | **2.78E-03** | 8.27E-04 | 7.09E-01 | | 2.99E-02 |
| 10 | **4.81E-03** | 8.27E-04 | **1.04E-03** | 5.85E-02 | 6.19E-03 |
| 11 | | **7.47E-07** | **6.51E-05** | | 2.60E-03 |
| 12 | | 5.93E-07 | **1.63E-05** | | 8.80E-05 |
| 13 | 8.42E-02 | 1.21E-05 | **1.04E-03** | 4.55E-02 | 3.87E-03 |
| 14 | | 9.15E-05 | 6.72E-02 | | **5.05E-04** |
| 15 | | **9.55E-09** | 3.46E-05 | 1.20E-01 | 2.04E-04 |
| 16 | **7.00E-06** | 2.37E-06 | 1.05E-03 | | **5.00E-06** |
| 17 | | 3.21E-01 | 1.18E-05 | **2.05E-02** | 1.03E-01 |
| 18 | | | | | 1.89E-01 |
| 19 | | 2.72E-03 | 1.71E-02 | | 1.52E-01 |
| 20 | **1.40E-03** | **5.37E-09** | 6.24E-02 | **5.28E-03** | **2.04E-04** |
| 21 | | 7.35E-03 | 4.18E-03 | | **3.24E-04** |
| 22 | 1.83E-02 | 1.73E-15 | 1.22E-07 | **1.48E-02** | **1.70E-05** |
| 23 | **2.78E-03** | | 1.24E-02 | | 8.63E-02 |

**Supplemental Figure 1. Cell- and tissue-level heatmap examples by cancer type**

Unprocessed portions of BRCA, LUAD, LUSC, and SKCM H&E-stained slides alongside corresponding overlays (heatmaps) of cell- and tissue-type predictions. Slide regions are classified into tissue types: cancer tissue (red), cancer-associated stroma (orange), necrosis (black), or normal (transparent). Pixels in cancer tissue or cancer-associated stroma areas are classified into cell types: lymphocyte (green), plasma cell (lime), fibroblast (orange), macrophage (aqua), cancer cell (red), or background (transparent). Examples for STAD are shown in Figure 1c.

### BRCA

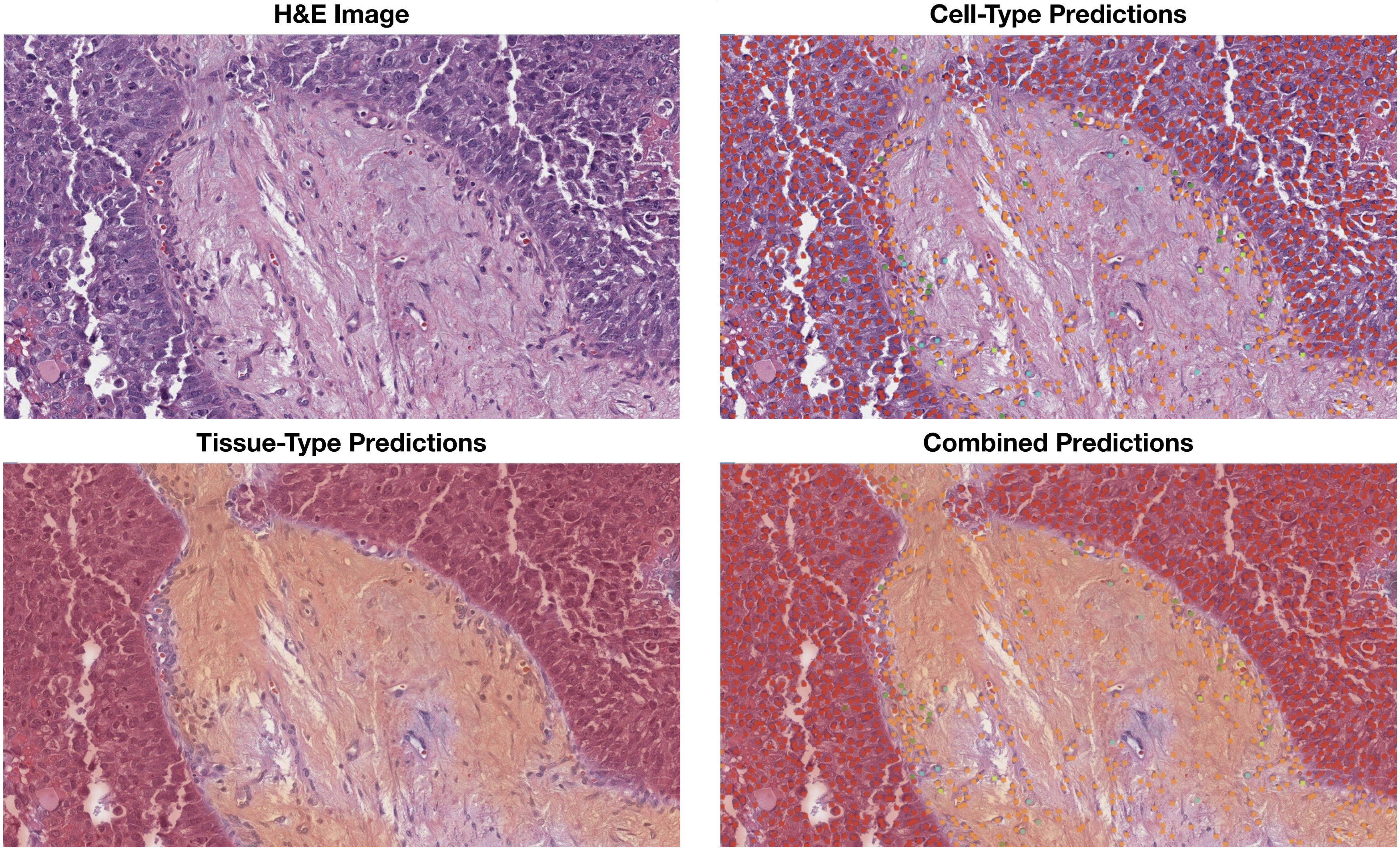
H&E Image
Cell-Type Predictions
Tissue-Type Predictions
Combined Predictions

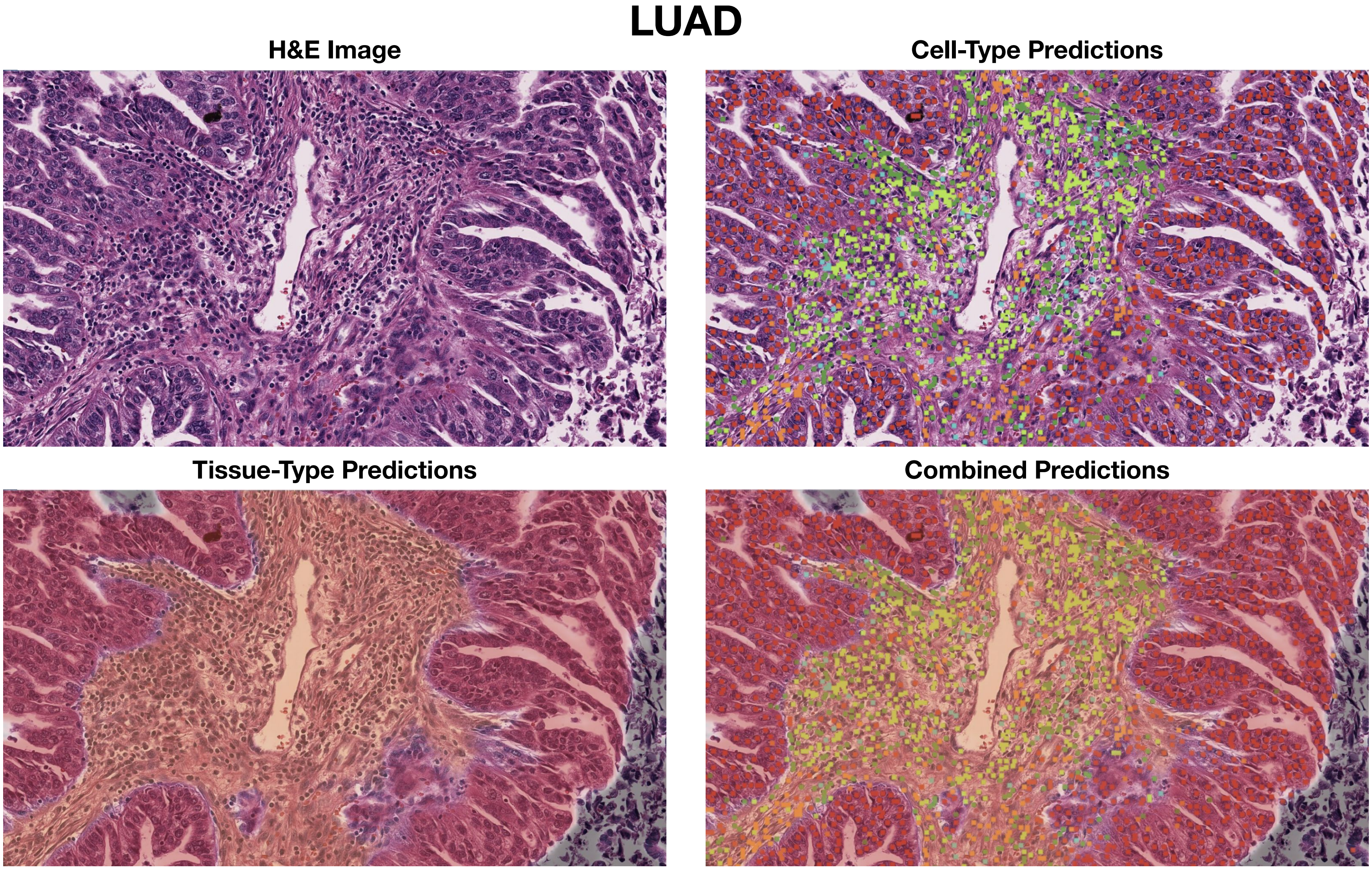
LUAD
H&E Image
Cell-Type Predictions
Tissue-Type Predictions
Combined Predictions

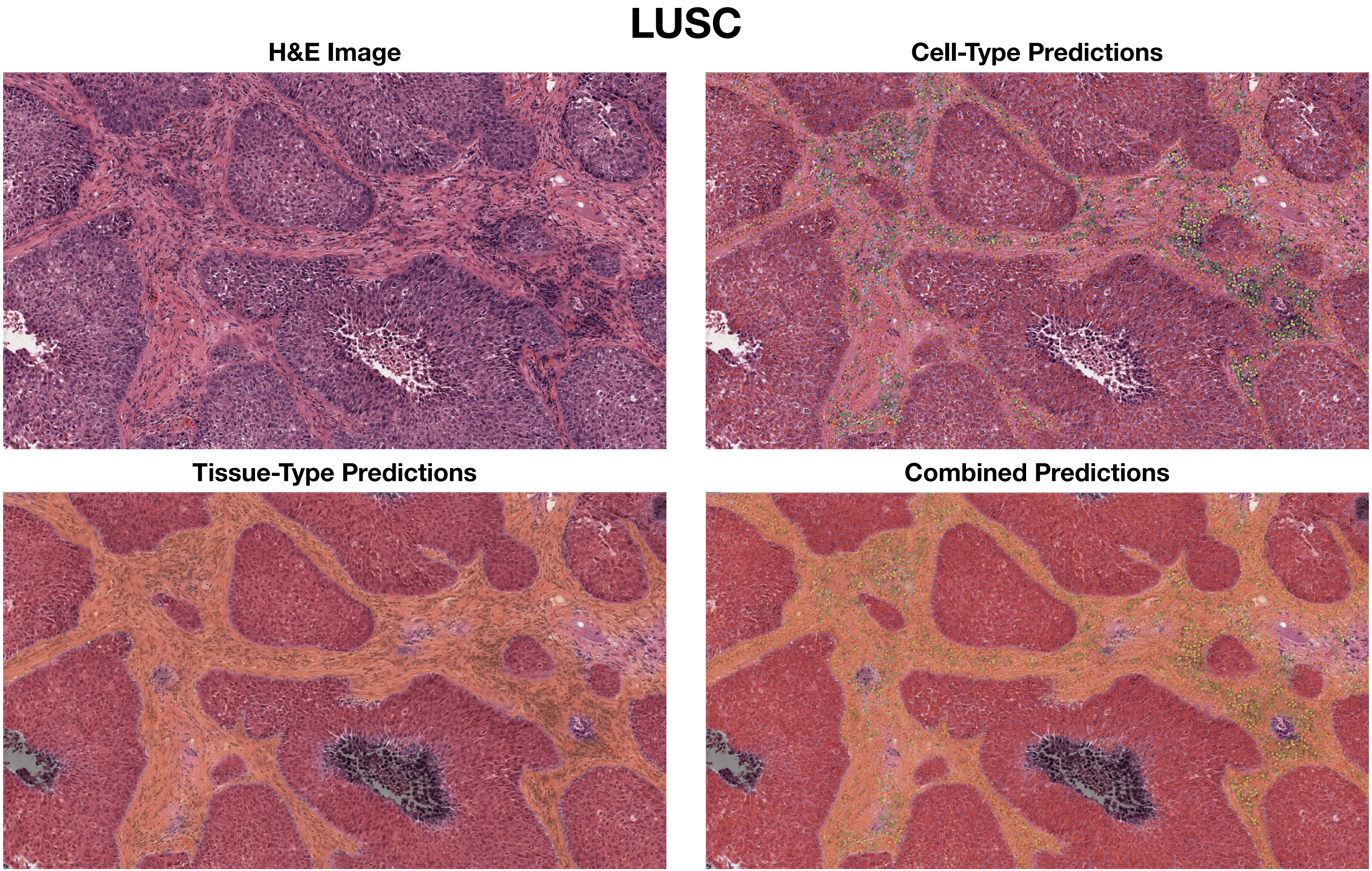
LUSC
H&E Image
Cell-Type Predictions
Tissue-Type Predictions
Combined Predictions

### SKCM

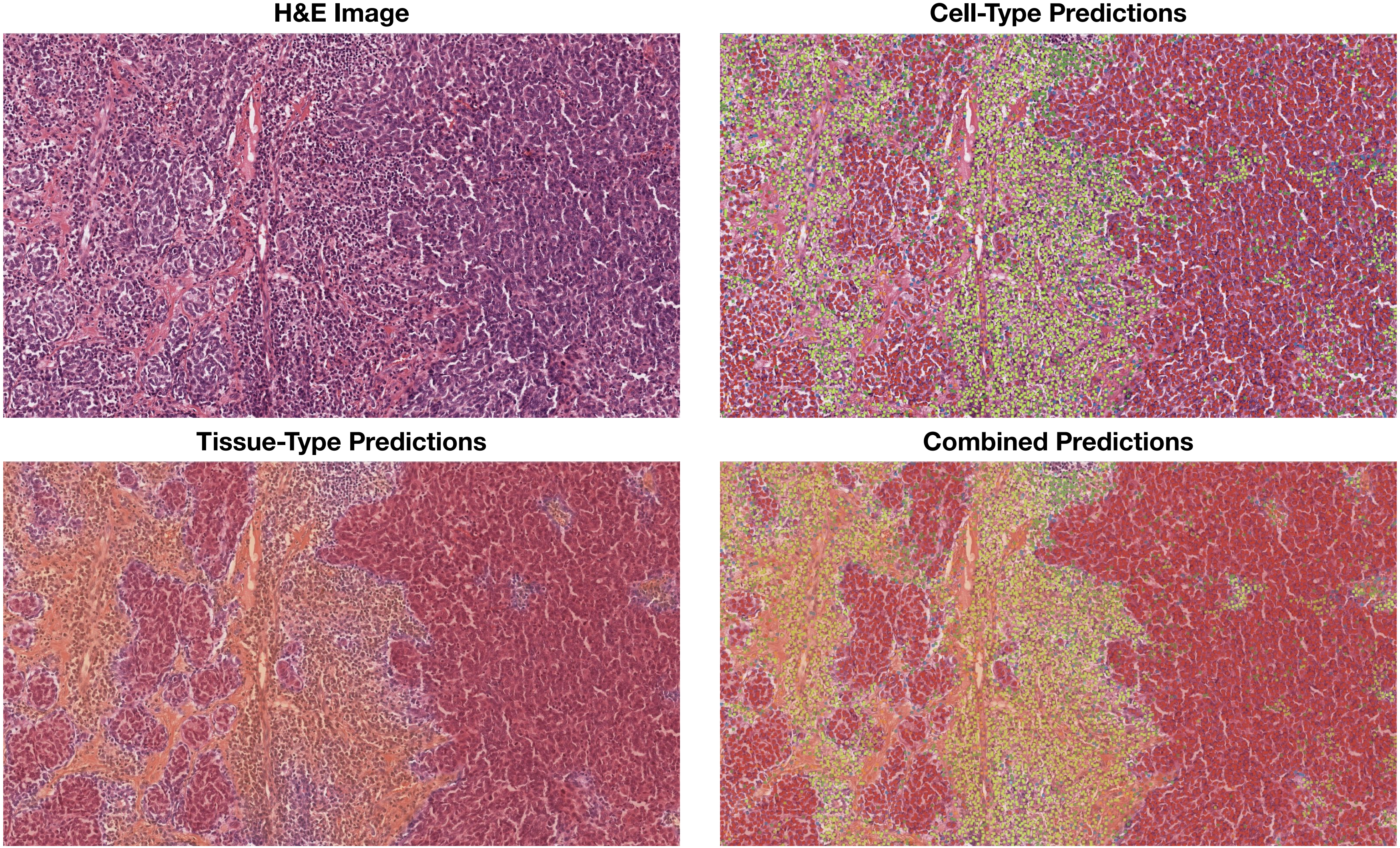

H&E Image
Cell-Type Predictions
Tissue-Type Predictions
Combined Predictions

**Supplemental Figure 2. Validation of HIFs against molecular deconvolution of immune cell contributions**

Comparison of leukocyte fraction, plasma cell fraction, and lymphocyte fraction estimates from our 2-micron resolution cell-type model predictions (x-axis) and CIBERSORT RNA-Seq deconvolutions (y-axis) across all patient samples. All measured Spearman correlations (0.40-0.55) are positive and consistent with informative image-based features. Potential sources of disagreement include RNA contributions from tumor, fibroblast, and other non-immune cells, discrepancies between 3D bulk samples and 2D tissue slices, tissue sampling variability, and RNA-Seq measurement variability.

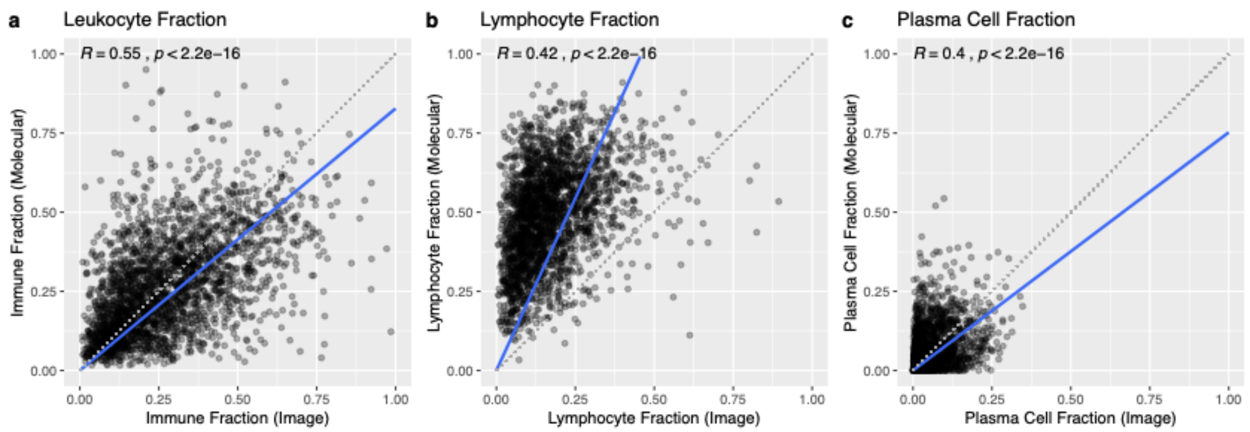

**Supplemental Figure 3. Gaussian mixture model thresholding**

Histogram of (a) PD-1 expression, (b) PD-L1 expression, (c) CTLA-4 expression, (d) HRD score, and (e) TIGIT expression broken down by cancer type. Continuous immune checkpoint protein expression and HRD scores were binarized to high versus low classes using gaussian mixture model (GMM) clustering with unequal variance. The binary thresholds annotated below were defined as the intersection of the empirical densities between the two GMM-defined clusters. The number and percentage of positive labels per outcome are shown.

**a)**

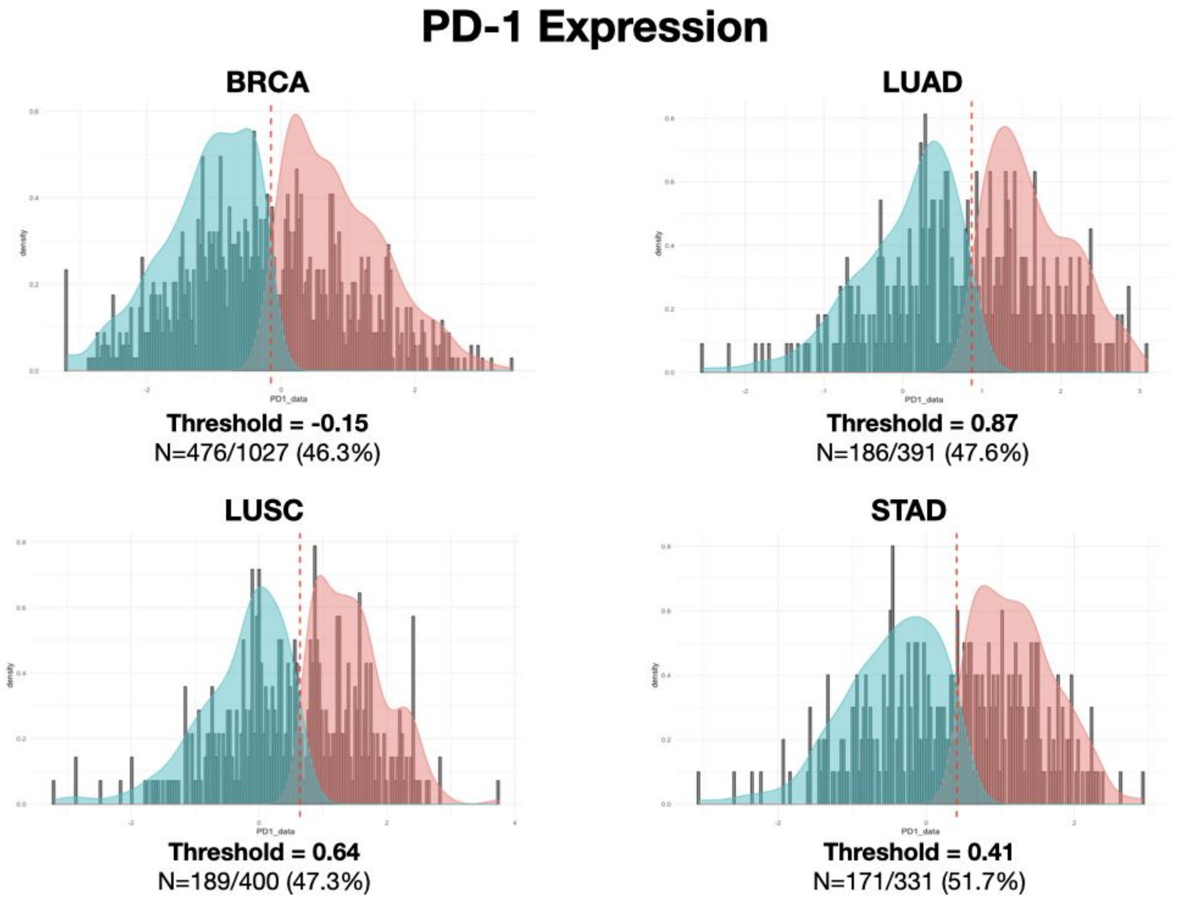

b)

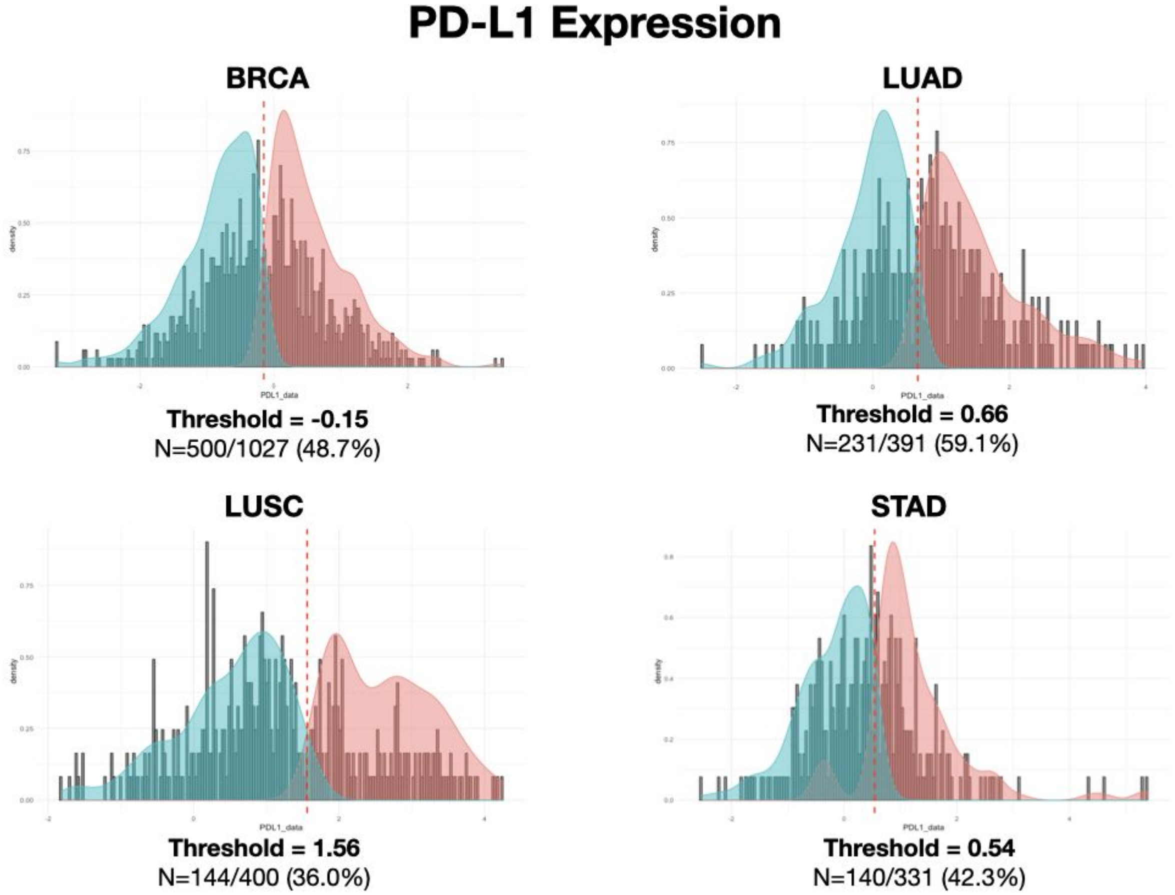
PD-L1 Expression
BRCA
Threshold = -0.15
N=500/1027 (48.7%)
LUAD
Threshold = 0.66
N=231/391 (59.1%)
LUSC
Threshold = 1.56
N=144/400 (36.0%)
STAD
Threshold = 0.54
N=140/331 (42.3%)

**c)**

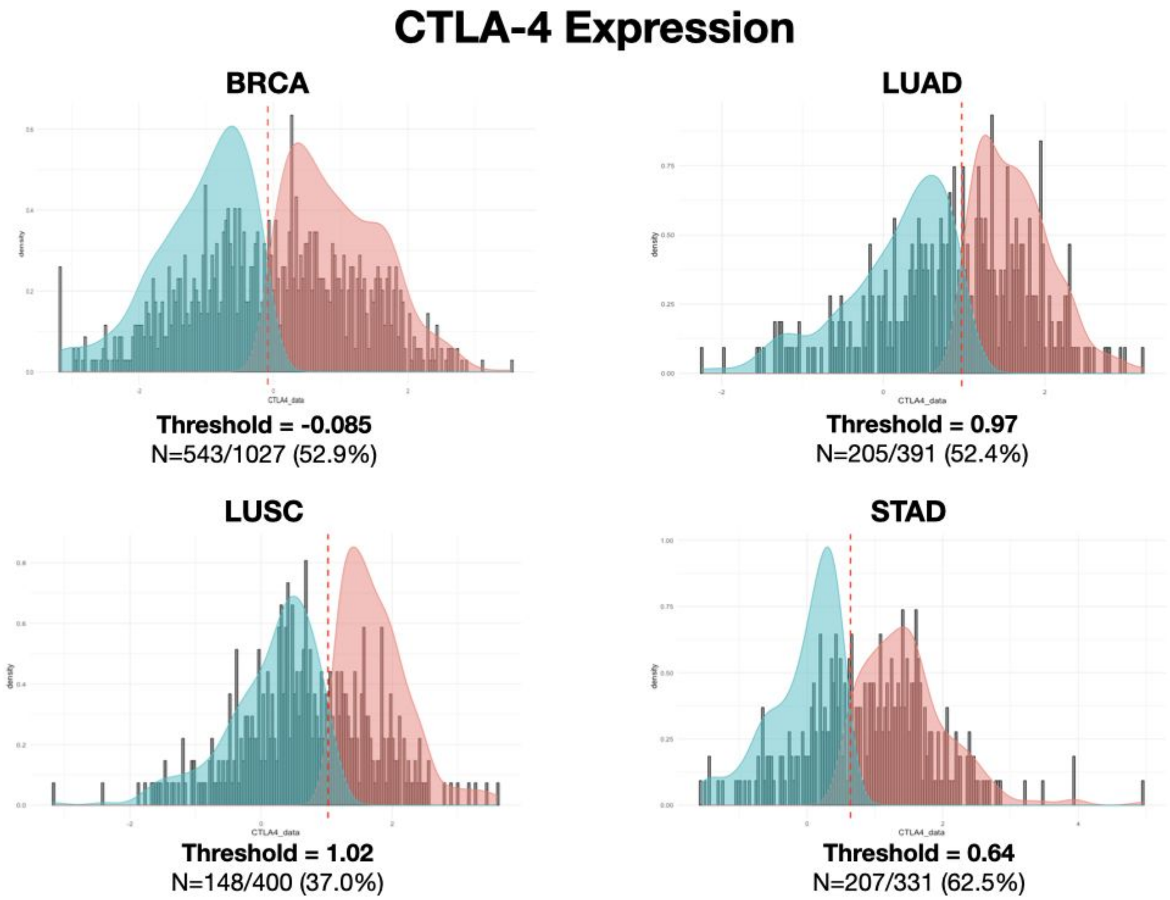
CTLA-4 Expression
BRCA
Threshold = -0.085
N=543/1027 (52.9%)
LUAD
Threshold = 0.97
N=205/391 (52.4%)
LUSC
Threshold = 1.02
N=148/400 (37.0%)
STAD
Threshold = 0.64
N=207/331 (62.5%)

**d)**

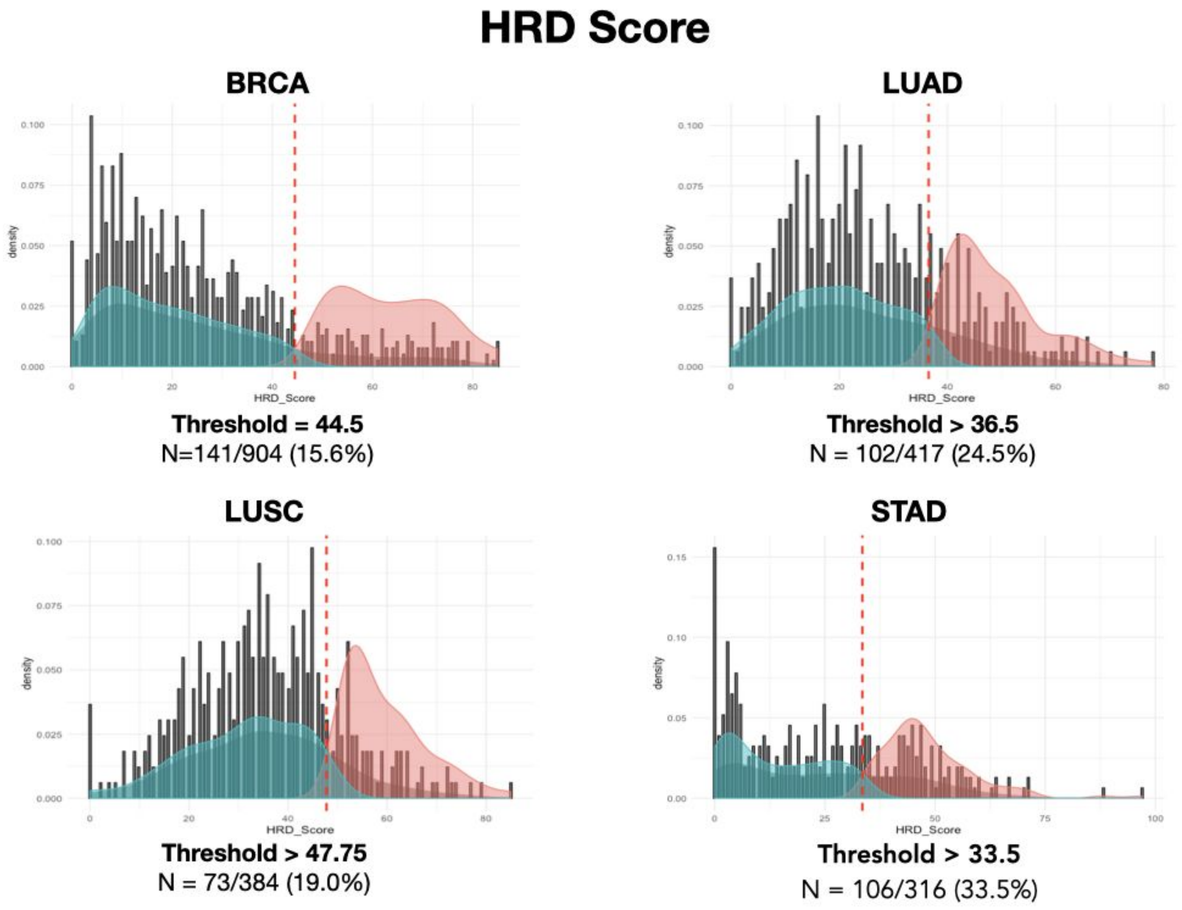
HRD Score
BRCA
Threshold = 44.5
N=141/904 (15.6%)
LUAD
Threshold > 36.5
N = 102/417 (24.5%)
LUSC
Threshold > 47.75
N = 73/384 (19.0%)
STAD
Threshold > 33.5
N = 106/316 (33.5%)
density
HRD_Score

**e)**

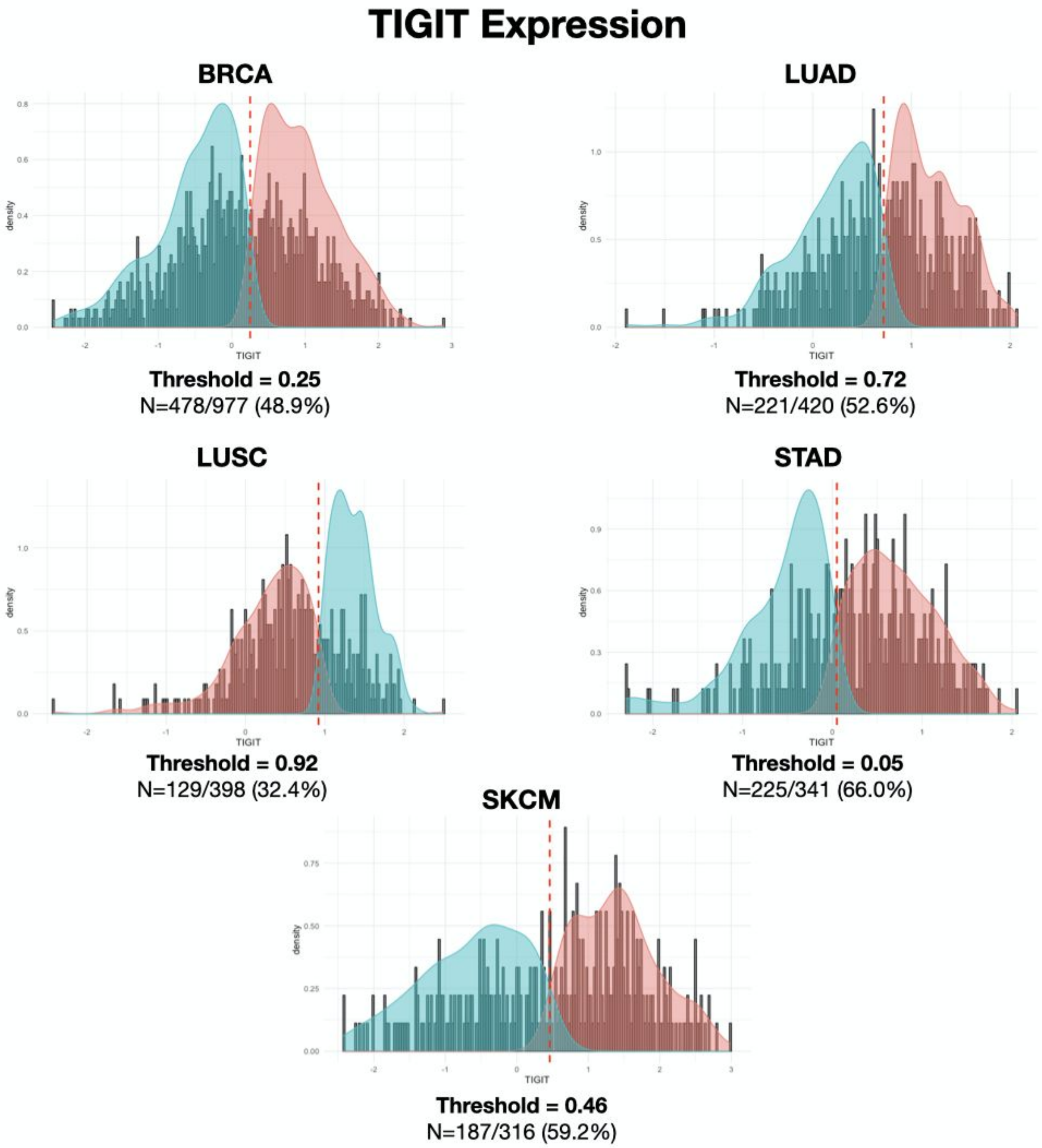
TIGIT Expression
BRCA
Threshold = 0.25
N=478/977 (48.9%)
LUAD
Threshold = 0.72
N=221/420 (52.6%)
LUSC
Threshold = 0.92
N=129/398 (32.4%)
STAD
Threshold = 0.05
N=225/341 (66.0%)
SKCM
Threshold = 0.46
N=187/316 (59.2%)
density
TIGIT

**Supplemental Figure 4. Precision-recall curves from prediction of molecular phenotypes using HIFs**

Precision-recall (PR) curves for hold-out predictions, colored by cancer type. A random classifier corresponds to an AUPRC equal to the percentage of positive labels for a given prediction task (i.e. a horizontal PR curve drawn at the said percentage).

Pan-cancer in TIGIT predictions encompasses all five cancer types. Pan-cancer for the remainder of prediction tasks excluded SKCM due to insufficient outcome labels.

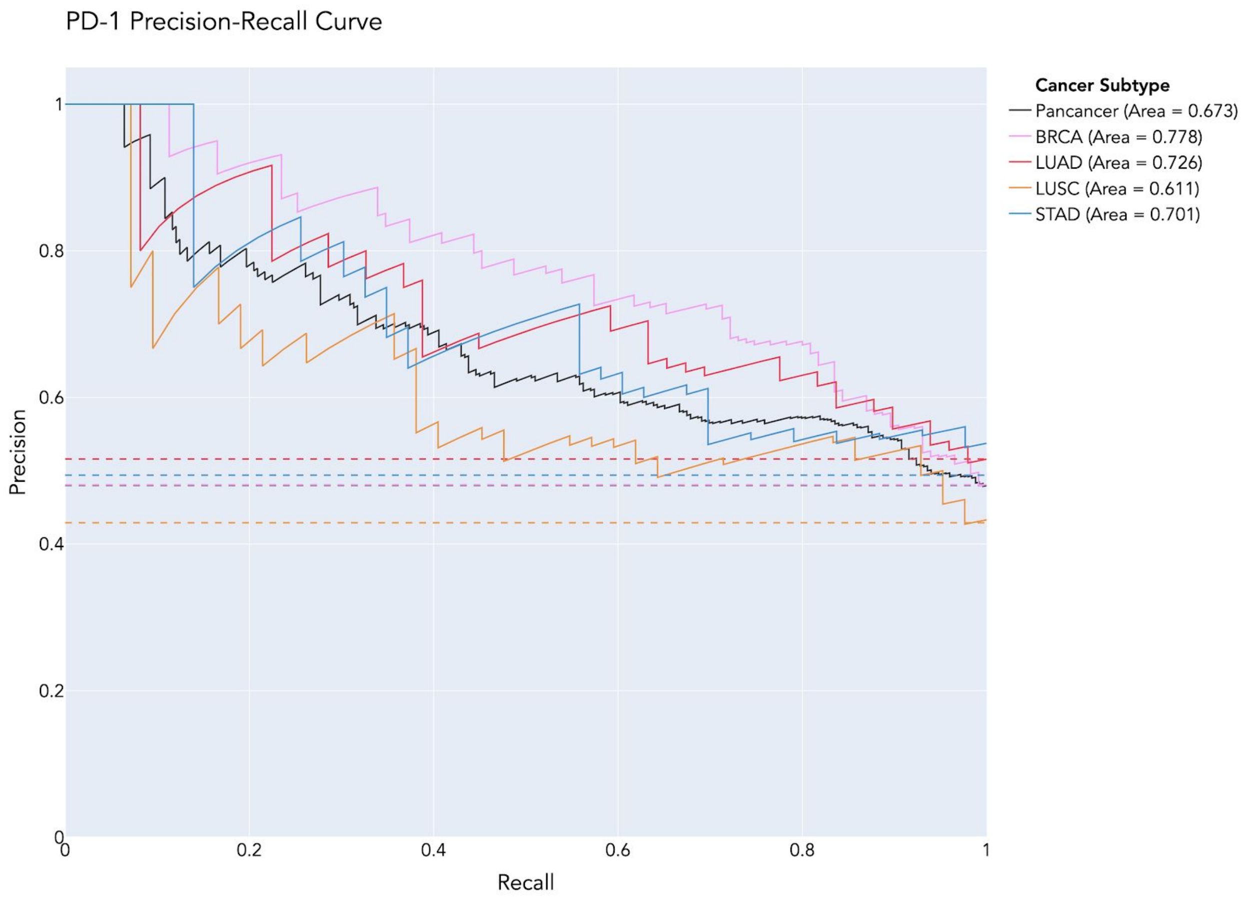

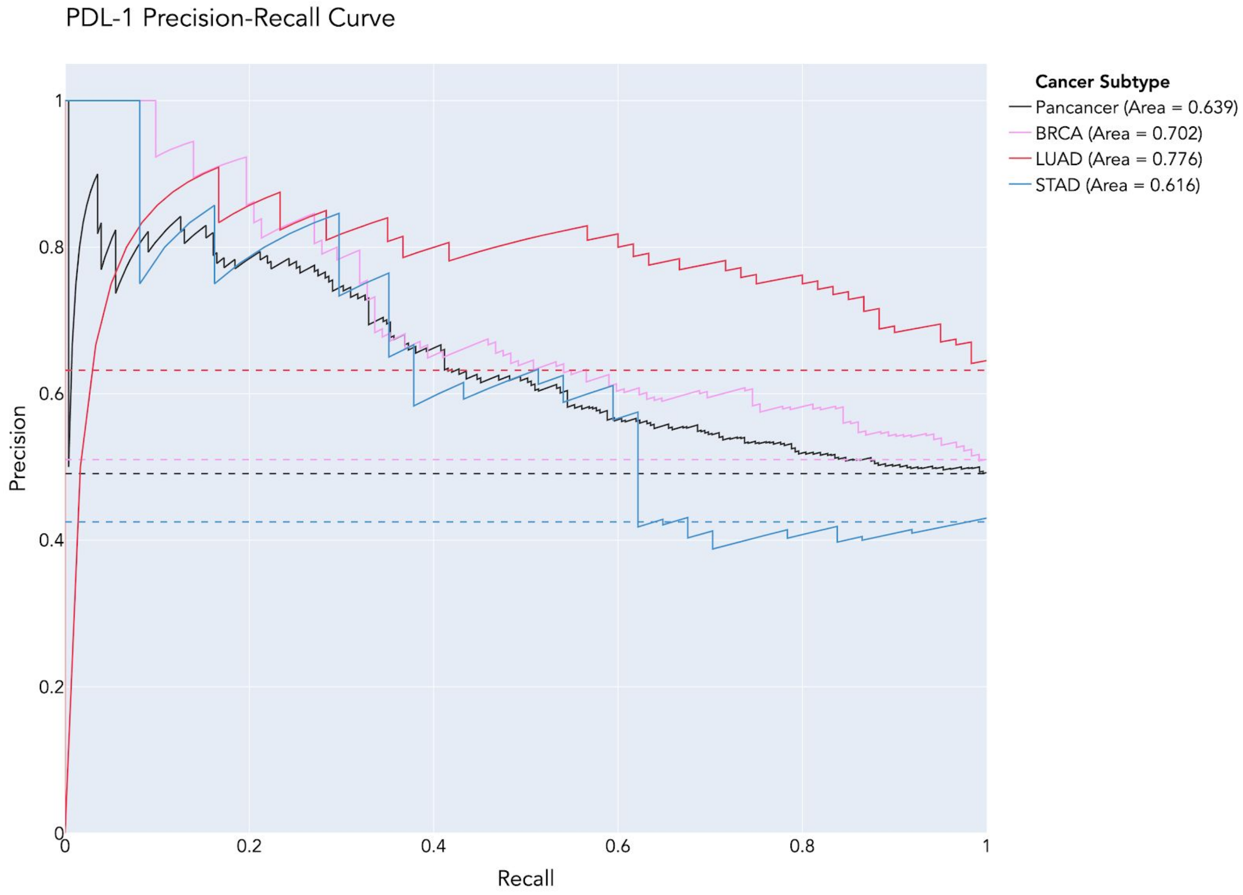
PDL-1 Precision-Recall Curve
Precision
Recall
0
0.2
0.4
0.6
0.8
1
Cancer Subtype
Pancancer (Area = 0.639)
BRCA (Area = 0.702)
LUAD (Area = 0.776)
STAD (Area = 0.616)

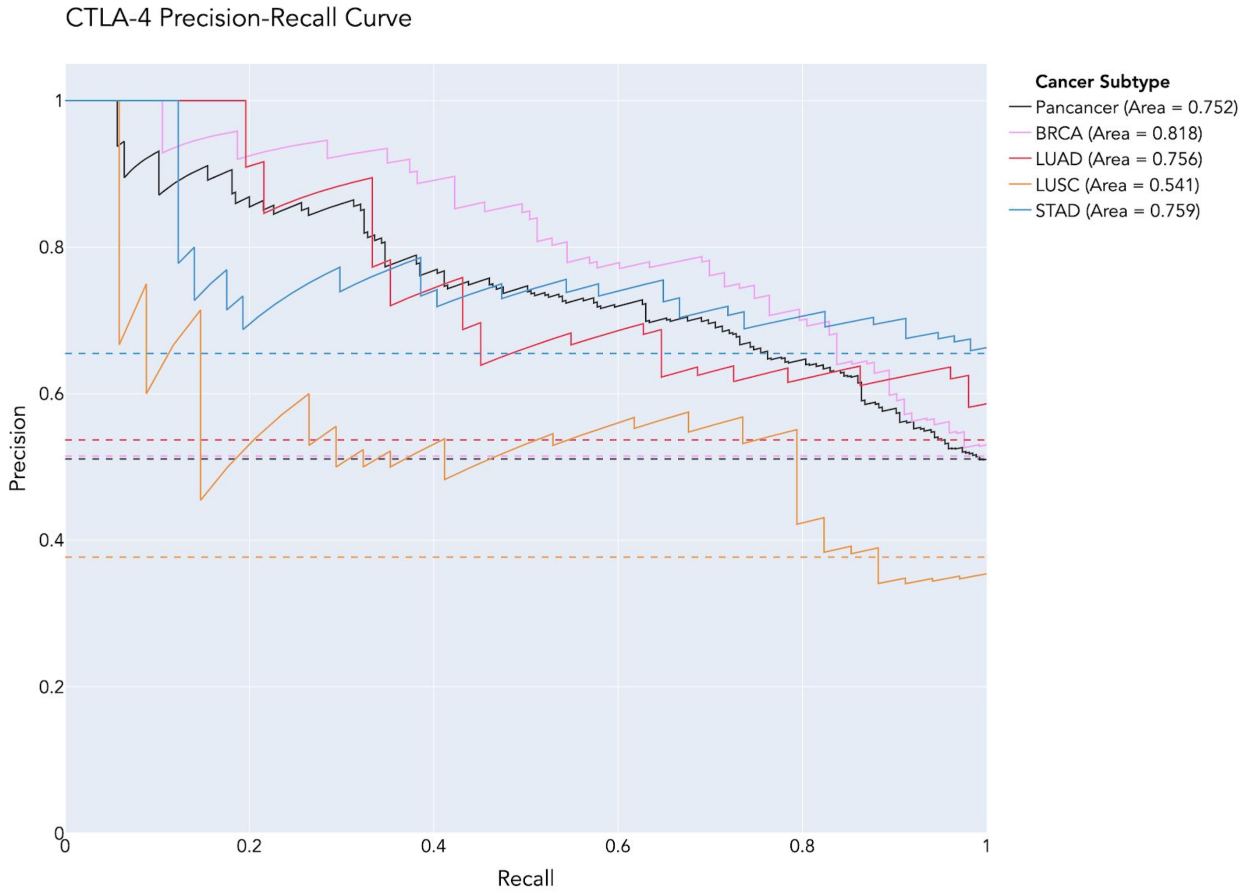
CTLA-4 Precision-Recall Curve
Cancer Subtype
Pancancer (Area = 0.752)
BRCA (Area = 0.818)
LUAD (Area = 0.756)
LUSC (Area = 0.541)
STAD (Area = 0.759)
Precision
Recall
0
0.2
0.4
0.6
0.8
1

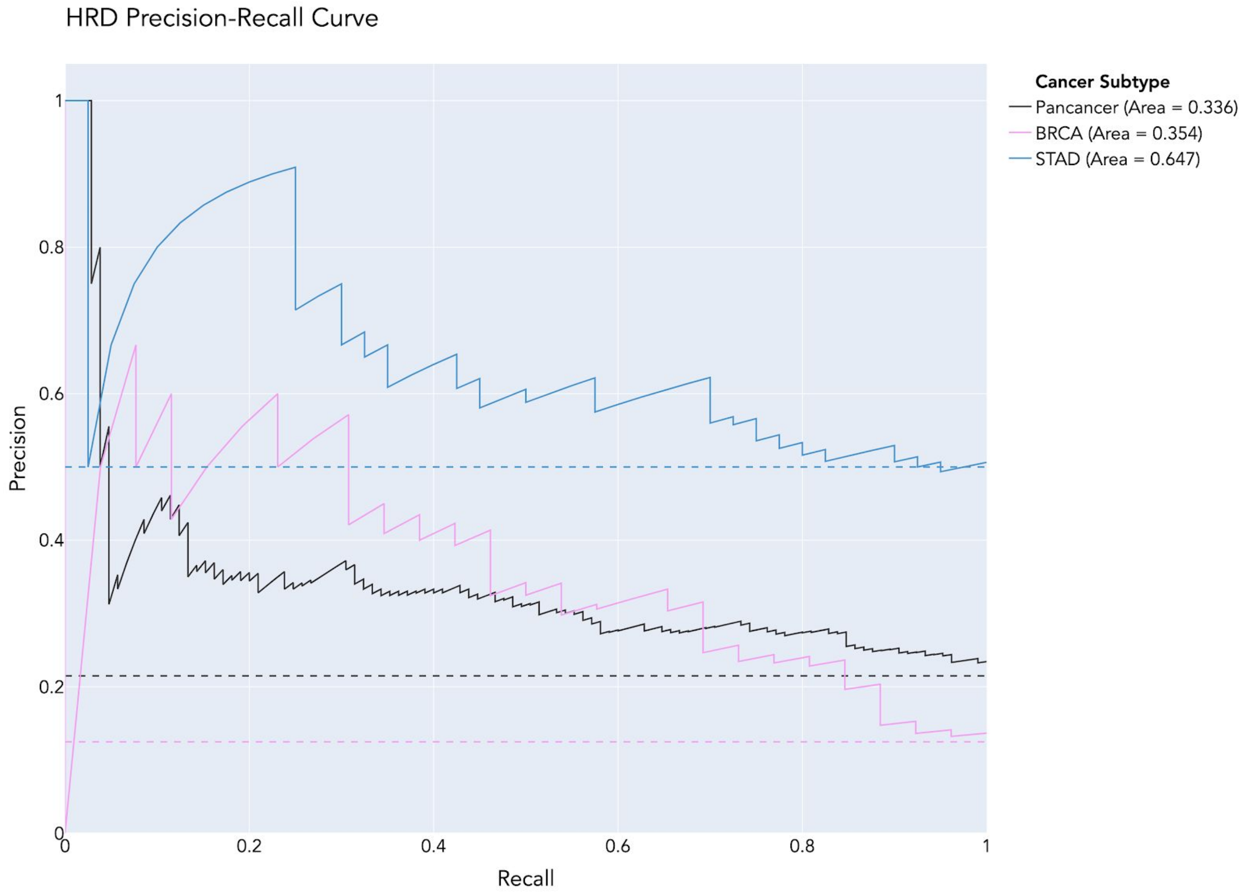

HRD Precision-Recall Curve
Cancer Subtype
Pancancer (Area = 0.336)
BRCA (Area = 0.354)
STAD (Area = 0.647)
Precision
Recall
0
0.2
0.4
0.6
0.8
1

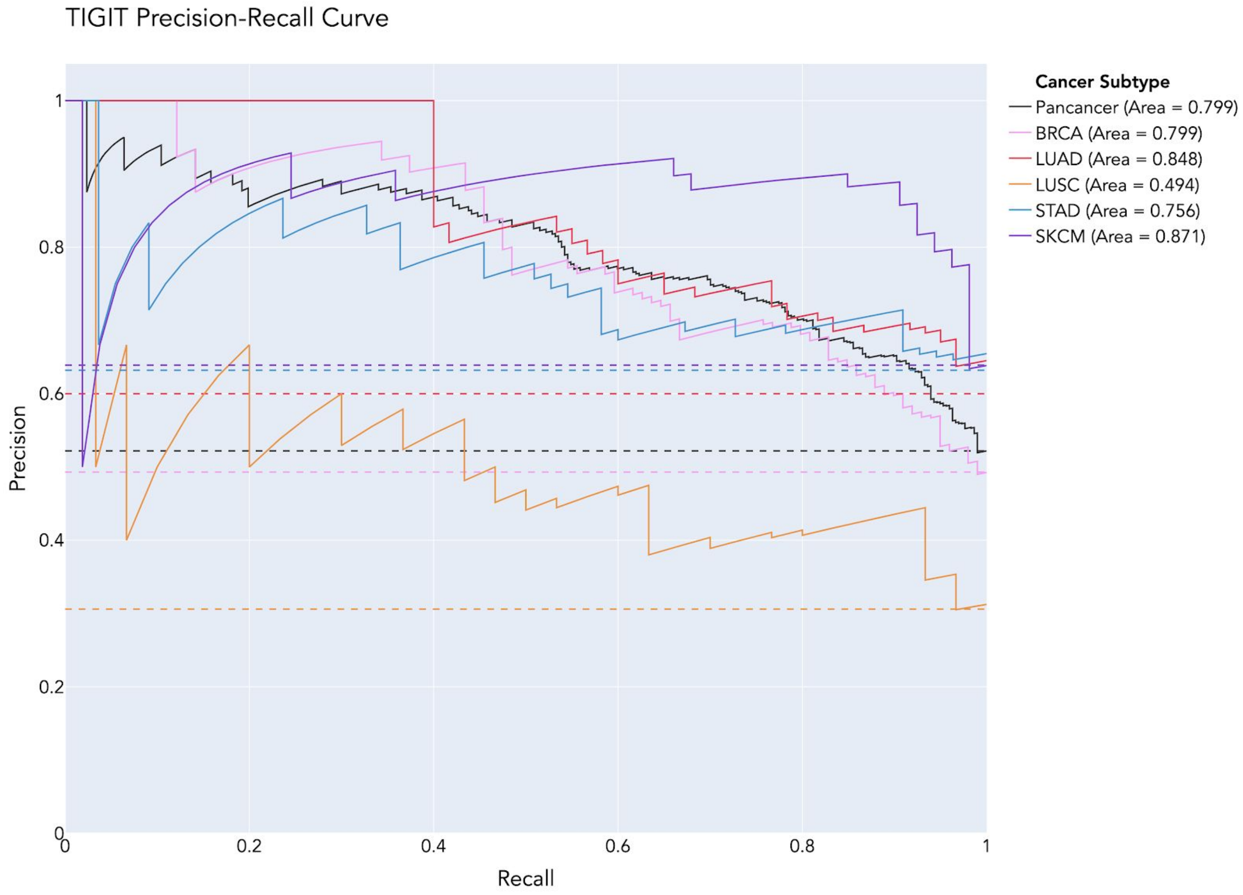

TIGIT Precision-Recall Curve
Cancer Subtype
Pancancer (Area = 0.799)
BRCA (Area = 0.799)
LUAD (Area = 0.848)
LUSC (Area = 0.494)
STAD (Area = 0.756)
SKCM (Area = 0.871)
Precision
Recall
0
0.2
0.4
0.6
0.8
1

**Supplemental Figure 5. Predictive HIF clusters for cancer type-specific models**
Boxplots of the top five most predictive HIF clusters (defined per cancer type) for predicting each molecular phenotype across cancer types and pan-cancer. Clusters are ranked by the maximum absolute beta across HIFs in a given cluster. Betas are computed per HIF as the average across the three models incorporated into the final ensemble evaluated on the hold-out set. Each boxplot highlights the median and interquartile range for HIF betas in each cluster. Each cluster is labeled with a representative HIF corresponding to the maximum absolute beta value. In cases in which that HIF is difficult to interpret, a more interpretable HIF within a five-fold difference of the maximum absolute beta is presented as indicated by a black asterisk. As the absolute value of beta values was used for ranking, HIFs with negative beta values are denoted by a red asterisk. For prediction tasks in which less than five HIF clusters have non-zero betas, only non-zero clusters are presented. Multiple predictive HIFs are visualized with overlaid cell or tissue-type heatmaps in Figure 3. Across all HIFs, tumor regions include cancer tissue (CT), cancer-associated stroma (CAS), and a combined CT+CAS.

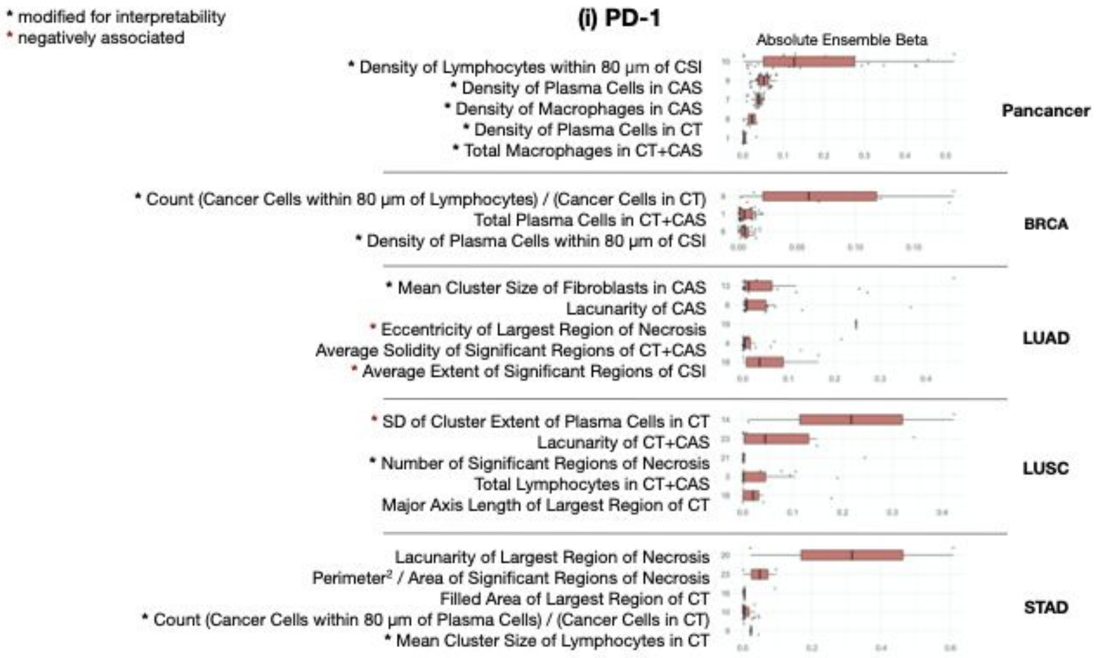

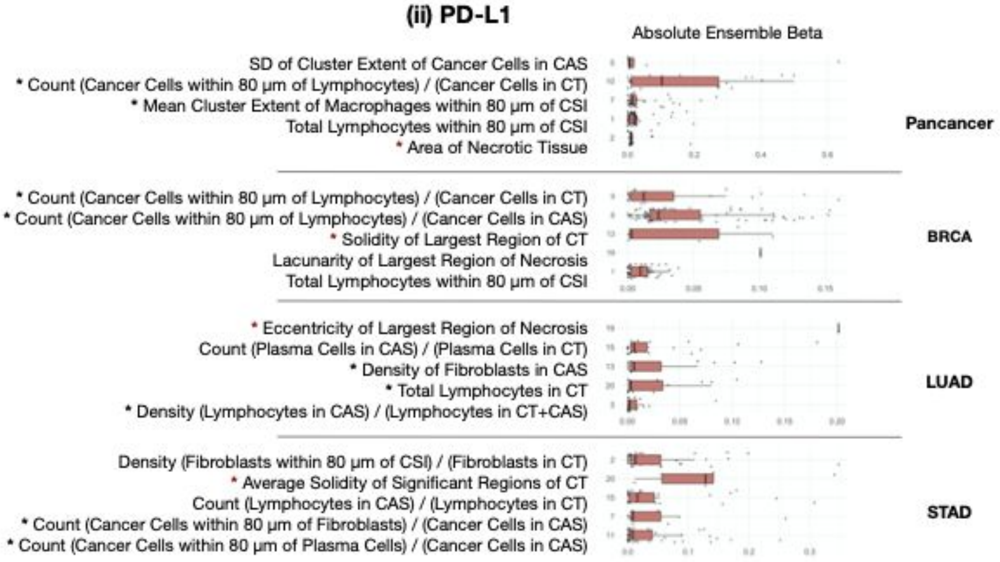
(ii) PD-L1
Absolute Ensemble Beta
SD of Cluster Extent of Cancer Cells in CAS
* Count (Cancer Cells within 80 µm of Lymphocytes) / (Cancer Cells in CT)
* Mean Cluster Extent of Macrophages within 80 µm of CSI
Total Lymphocytes within 80 µm of CSI
* Area of Necrotic Tissue
Pancancer
* Count (Cancer Cells within 80 µm of Lymphocytes) / (Cancer Cells in CT)
* Count (Cancer Cells within 80 µm of Lymphocytes) / (Cancer Cells in CAS)
* Solidity of Largest Region of CT
Lacunarity of Largest Region of Necrosis
Total Lymphocytes within 80 µm of CSI
BRCA
* Eccentricity of Largest Region of Necrosis
Count (Plasma Cells in CAS) / (Plasma Cells in CT)
* Density of Fibroblasts in CAS
* Total Lymphocytes in CT
* Density (Lymphocytes in CAS) / (Lymphocytes in CT+CAS)
LUAD
Density (Fibroblasts within 80 µm of CSI) / (Fibroblasts in CT)
* Average Solidity of Significant Regions of CT
Count (Lymphocytes in CAS) / (Lymphocytes in CT)
* Count (Cancer Cells within 80 µm of Fibroblasts) / (Cancer Cells in CAS)
* Count (Cancer Cells within 80 µm of Plasma Cells) / (Cancer Cells in CAS)
STAD

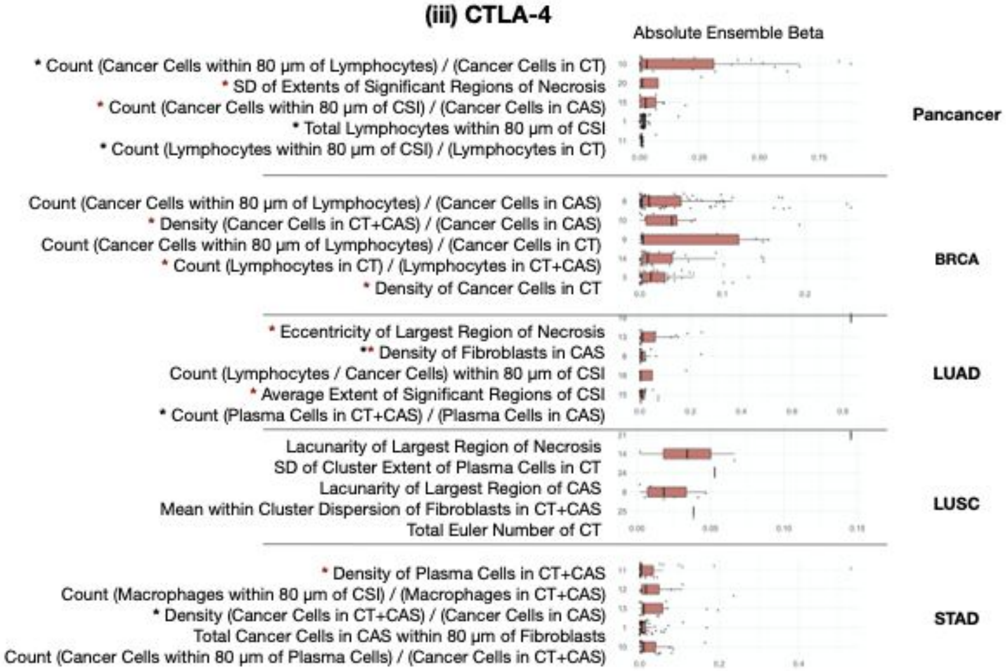
(iii) CTLA-4
Absolute Ensemble Beta
* Count (Cancer Cells within 80 µm of Lymphocytes) / (Cancer Cells in CT)
* SD of Extents of Significant Regions of Necrosis
* Count (Cancer Cells within 80 µm of CSI) / (Cancer Cells in CAS)
* Total Lymphocytes within 80 µm of CSI
* Count (Lymphocytes within 80 µm of CSI) / (Lymphocytes in CT)
Pancancer
Count (Cancer Cells within 80 µm of Lymphocytes) / (Cancer Cells in CAS)
* Density (Cancer Cells in CT+CAS) / (Cancer Cells in CAS)
Count (Cancer Cells within 80 µm of Lymphocytes) / (Cancer Cells in CT)
* Count (Lymphocytes in CT) / (Lymphocytes in CT+CAS)
* Density of Cancer Cells in CT
BRCA
* Eccentricity of Largest Region of Necrosis
** Density of Fibroblasts in CAS
Count (Lymphocytes / Cancer Cells) within 80 µm of CSI
* Average Extent of Significant Regions of CSI
* Count (Plasma Cells in CT+CAS) / (Plasma Cells in CAS)
LUAD
Lacunarity of Largest Region of Necrosis
SD of Cluster Extent of Plasma Cells in CT
Lacunarity of Largest Region of CAS
Mean within Cluster Dispersion of Fibroblasts in CT+CAS
Total Euler Number of CT
LUSC
* Density of Plasma Cells in CT+CAS
Count (Macrophages within 80 µm of CSI) / (Macrophages in CT+CAS)
* Density (Cancer Cells in CT+CAS) / (Cancer Cells in CAS)
Total Cancer Cells in CAS within 80 µm of Fibroblasts
Count (Cancer Cells within 80 µm of Plasma Cells) / (Cancer Cells in CT+CAS)
STAD

#### (iv) HRD

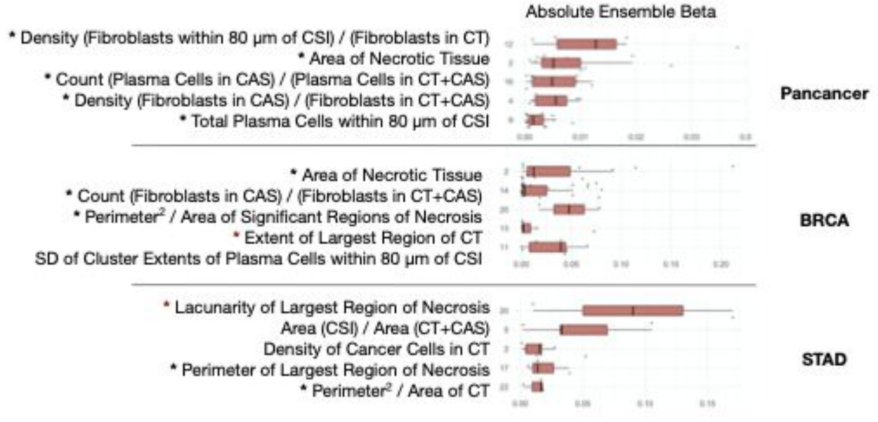


#### (v) TIGIT

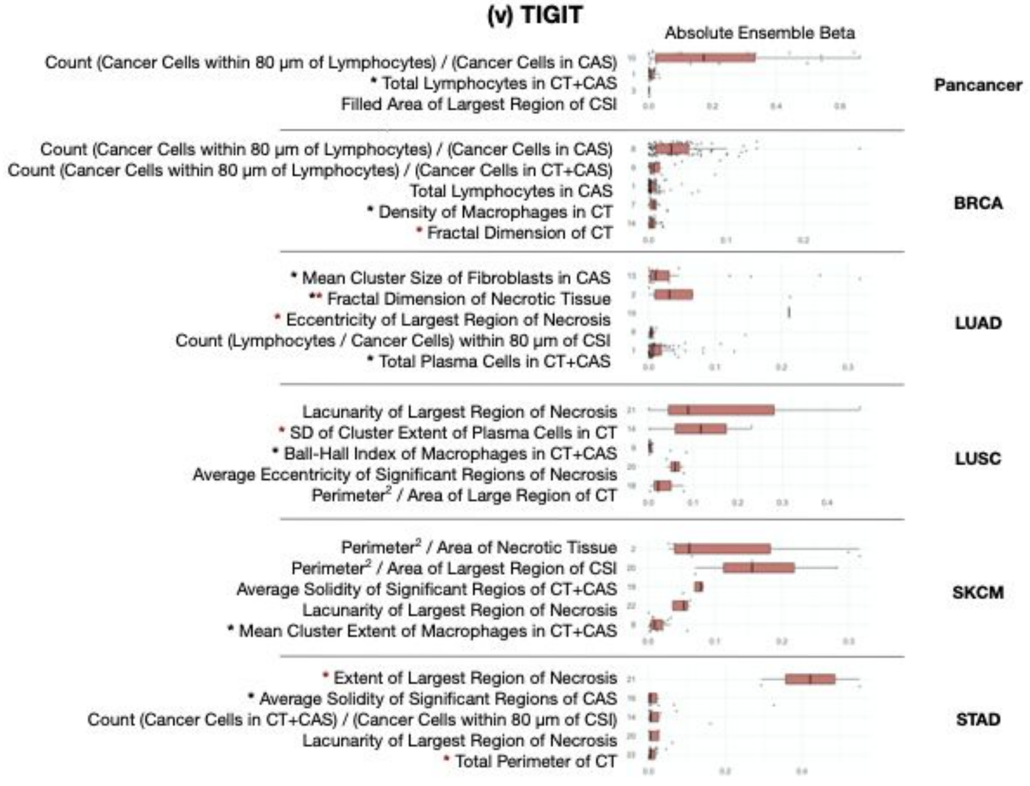

**Supplemental Figure 6. Molecular phenotype concordance**
Clustered heatmaps of concordance metrics between the five predicted molecular phenotypes computed on 1,893 patient samples across all five cancer types. In (a), Pearson correlations were computed between phenotypes in continuous form. In (b), the percentage agreement was computed between phenotypes after conversion into binary labels. We observe a strongly correlated cluster among the four immune checkpoint protein phenotypes (PD-1, PD-L1, CTLA-4, and TIGIT). HRD score appears relatively un-correlated with immune phenotypes.

**a)**

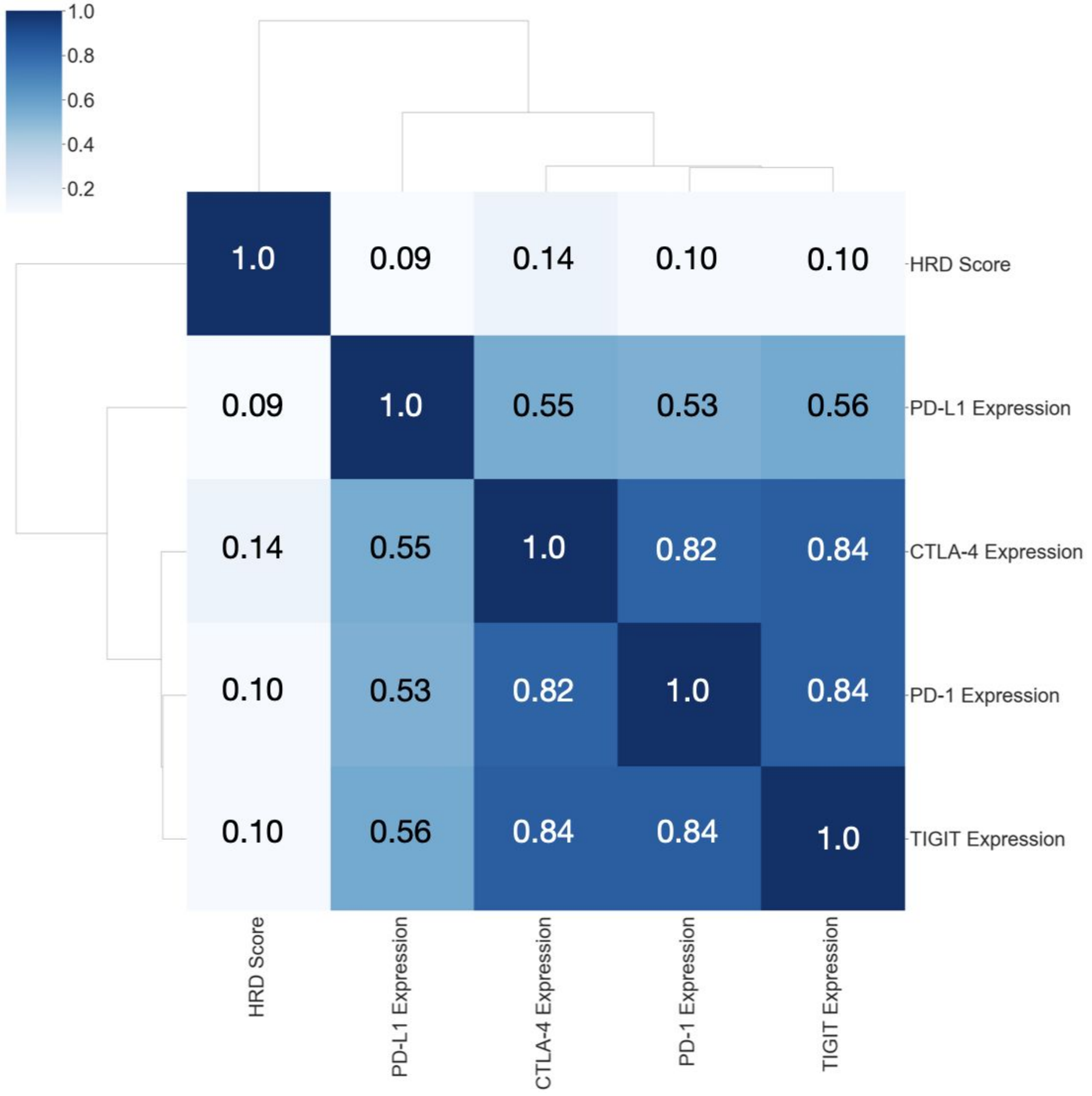

b)

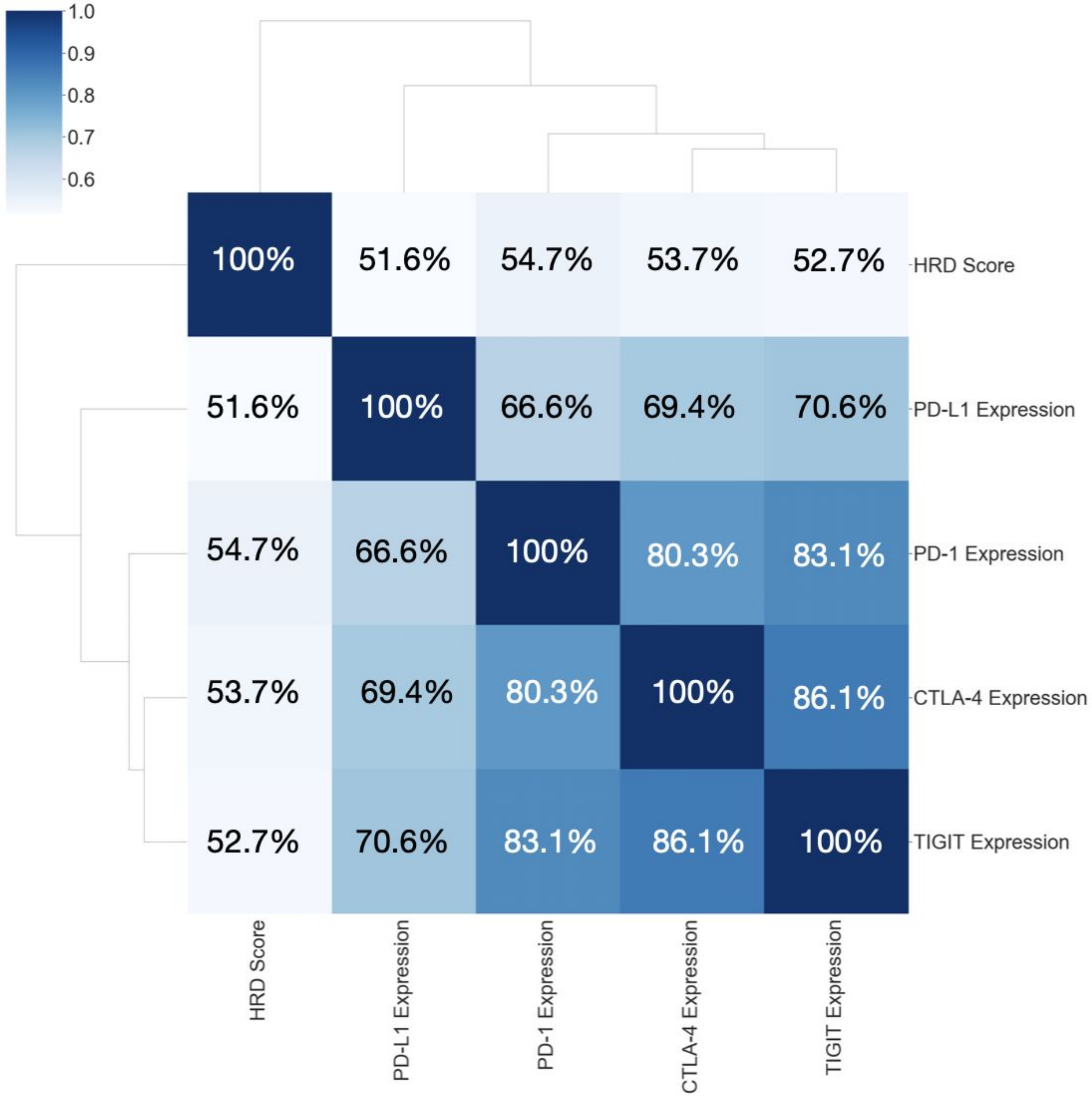
1.0
0.9
0.8
0.7
0.6
100%
51.6%
54.7%
53.7%
52.7%
66.6%
69.4%
70.6%
80.3%
83.1%
86.1%
HRD Score
PD-L1 Expression
PD-1 Expression
CTLA-4 Expression
TIGIT Expression
